# Time-series RNA-Seq and data-driven network inference unveil dynamics of cell activation, survival and crosstalk in Chronic Lymphocytic Leukaemia *in vitro* models

**DOI:** 10.1101/2025.04.20.649300

**Authors:** Malvina Marku, Hugo Chenel, Julie Bordenave, Marcelo Hurtado, Marcin Domagala, Flavien Raynal, Mary Poupot, Loïc Ysebaert, Andrei Zinovyev, Vera Pancaldi

**Author notes:** These authors have contributed equally (Malvina Marku, Hugo Chenel). Corresponding author: Vera Pancaldi, Malvina Marku.

## Abstract

How do cancer cells respond to their environment, and what are the key regulators behind their behaviour? While immune cell reprogramming in the tumour microenvironment (TME) has been extensively studied, the dynamic regulatory changes within cancer cells in response to interactions with immune cells remain poorly understood. In Chronic Lymphocytic Leukaemia (CLL), this knowledge gap limits our ability to fully grasp the disease progression and to design effective, personalised interventions. To tackle this, we combine time-series transcriptomics with data-driven gene regulatory network (GRN) inference to uncover the temporal regulatory mechanisms driving CLL cell behaviour within a reconstituted *in vitro* TME. Using cultures of peripheral blood from CLL patients or of purified patient-derived CLL cells, we profile gene expression across five time points spanning 14 days under these experimental conditions. By inferring GRNs from transcription factor activity, we capture patient-specific and temporally resolved regulatory interactions that highlight how immune signals shape cancer cell phenotypic changes. Our network analysis reveals distinct gene modules associated with critical processes such as cytokine signalling, metabolic reprogramming and differentiation, hallmarks of immune-cancer cell interaction. Intriguingly, we found that while the presence of immune cells in the environment significantly alters CLL cell activation, their survival trajectories are predominantly governed by intrinsic features. This study not only offers mechanistic insights into how immune cell presence influences CLL cell fate but also presents a robust computational framework for integrating time-series transcriptomics with GRN inference, which can then be used to study the long-term behaviour of the CLL cells through dynamical modelling.

## Introduction

The tumour microenvironment (TME) is a complex ecosystem of interactions between immune and stromal cells, the extracellular matrix and cancer cells, occurring inside tumours from molecular to cellular scales. Studying its temporal evolution allows us to define cancer cells’ cellular behaviour and their response to external stimuli. In the presence of cancer cells, several immune cells, including T cells, macrophages and neutrophils, undergo cell-state transitions toward pro-tumoural phenotypes or exhaustion (Ando et al., 2020; Fridlender et al., 2009; Jiang et al., 2015; Masucci et al., 2019; Mishalian et al., 2013), compromising the action of the immune system. Cellular phenotypic shifts and state transitions of immune cells are well studied, both experimentally and computationally, while their impact on the dynamics of the TME remains an important subject of investigation. For example, it is widely studied that macrophages found in tumours, denoted as tumour-associated macrophages (TAMs), can be educated by the cancer cells to promote their resistance to immune attacks and therapies, favouring tumour growth (Mantovani et al., 2022; Noy and Pollard, 2014; Sheban et al., 2025). The activation of the pro-tumoral phenotype in TAMs leads to the secretion of several cytokines, such as CXCL12/13, IL-10, and IL-6/IL-8, which are reported to promote tumour development by protecting the cancer cells (Cassetta and Pollard, 2018).

In system biology, the cellular processes that determine cell state transitions can be represented as gene regulatory networks (hereafter, GRNs) in which nodes represent proteins, enzymes, chemokines, etc., while the connections represent the type (activation or inhibition) and direction of interaction (Marku et al., 2023). Network modelling can be used to perform *in silico* experiments to predict how internal or external perturbations can drive these state transitions and how they impact interactions between cancer and immune cells (Cacace et al., 2020; Kondratova et al., 2020; Marku et al., 2020). As a matter of fact, our understanding of the transition dynamics of cancer cell states in response to the interactions with immune cells remains partial, particularly considering the context-specificity or patient variability. Several mathematical models - in particular Boolean models - of cancer cells for various cancer types have been built (Chuang et al., 2015; Flobak et al., 2015, 2015; Folkesson et al., 2023; Montagud et al., 2022), particularly focusing on identifying novel drug targets, studying the synergistic effect of drug combinations, or treatment optimisation. However, the context-specificity of the tumour microenvironment in which these cells have been characterised remains hidden, unexplored, or generalised among different sources of information used for model building. In addition, these models have been mostly developed based on public databases of gene interactions or the literature, thus being focused on a restricted number of well-studied regulatory pathways or drug targets. Data-driven GRN inference has been an ongoing research topic for more than 25 years, leading to the development of numerous inference methods, employing several algorithms, and building on different types of molecular data (Delgado and Gómez-Vela, 2019; Huynh-Thu and Sanguinetti, 2019; Marku and Pancaldi, 2023; Mercatelli et al., 2020). Despite the remaining challenges in the field, GRN inference leveraging different types of ‘omics’ data and prior knowledge helps to characterise pathways, biological components, and interactions in a concrete cellular context, thus further increasing our understanding of underlying cellular mechanisms and opening the way for creating executable and predictive models.

In this work, we follow a data-driven approach to infer a GRN of primary Chronic Lymphocytic Leukaemia cells (hereafter, CLL) and explore their behaviour when interacting with other immune cells. CLL is a haematological malignancy, characterised by the accumulation of large quantities of CD19+/CD5+ B cancer cells in the bloodstream and bone marrow. However, it is within secondary lymphoid organs, such as the spleen and lymph nodes, that CLL cells can interact with a supportive TME that promotes their proliferation and drives disease progression (Burger, 2011; Hayden et al., 2012; Ponzoni et al., 2011). Unlike other cancer types, CLL cells rely on extrinsic cues for proliferation and survival, requiring migration to specialised niches known as proliferation centres. These niches are rich in T cells, stromal elements, and TAMs, referred to in CLL as Nurse-Like Cells (NLCs), which provide key survival signals (Filip et al., 2013; Fiorcari et al., 2021; Svanberg et al., 2021). *In vitro* studies have widely reported that NLCs are crucial in rescuing CLL cells from spontaneous apoptosis (Boissard et al., 2016; Domagala et al., 2022) and are important in attracting them to the proliferation centres (Filip et al., 2015; Kipps et al., 2017). T cells can also play a key role in CLL cell proliferation (Hoferkova et al., 2022). CLL cells proliferate near activated helper T cells, while their proliferation *in vitro* requires direct contact with pre-activated T cells (Hoferkova et al., 2022).

In addition, NLCs are known to secrete various cytokines and chemokines, which can impact T cells by promoting an immunosuppressive environment that may induce a shift of T cells into T helper cells. *In vitro* cultures of Peripheral Blood Mononuclear Cells (PBMC) have been key to understanding these dialogues in a simplified biological system in which we can easily collect data and apply specific conditions (Cadot et al., 2020).

Exploiting temporally resolved bulk RNAseq profiling of CLL cells cultured *in vitro* with and without other immune cells from patients’ peripheral blood, we perform an extensive analysis to understand and describe CLL cells’ dynamic phenotype in relation to their environment and study what determines their survival. Furthermore, we perform GRN inference to characterise gene interactions regulating the CLL cell phenotype in these conditions and investigate how different processes in the system are defined by the modular structure of the network. CLL is a cancer intricately tied to its environment. By adopting a data-driven approach and leveraging time series data from *in vitro* experiments, we aim to identify critical pathways involved in the interaction between CLL cells and immune cells within the TME, with potential relevance to solid tumours.

## Results

### Experimental setup of cellular cultures to generate transcriptomics time courses

To investigate the impact of immune cell presence on CLL cell phenotypic profiling and survival, we conduct *in vitro* experiments using patient-derived CLL cell cultures under two distinct conditions: **(1)** autologous cultures of PBMC from CLL patients (𝑛 = 3), and **(2)** monocultures of isolated CLL cells from the same patients (𝑛 = 3). The cultures are monitored over 14 days, and for each condition and each patient, two technical replicates are collected. In the autologous culture, we expect a low percentage of monocytes (typically between 2 and 10%), which differentiate into NLCs upon contact with the culture dish, becoming adherent while other cells remain in suspension. Flow cytometry analysis is performed by Fluorescence Assisted Cell Sorting (FACS) on the non-adherent cells at D1 of the autologous culture, showing a high percentage of CLL cells in the samples (97.69%, 90.07% and 97.91% in patient 1, 2 and 3 respectively), and much lower percentages of other lymphoid cells (T cells, Natural Killer cells) with proportions lower than 7% of the total (see Materials and Methods, and <u>Supplementary Materia</u>l). Similar results are also obtained from performing deconvolution analysis on the autologous cultures using CLL PBMC single-cell signatures (see Supplementary Material, Figure S.1). For both monoculture and autologous culture, cell counting and bulk RNA sequencing are performed on the non-adherent cells in the cultures at five time points (day 1, 4, 8, 11, and 14) (Figure 1 (a) and <u>Materials and Methods</u>). The selection of the time points is based on our past *in vitro* studies (Boissard et al., 2016; Verstraete et al., 2023), where we have monitored the formation of monocyte-derived NLCs upon co-culturing with CLL cells. We observe that monocyte-derived adherent macrophages begin transitioning to the NLC phenotype between D2 and D6, reaching their final state around D8 (as seen also in microscopy images, Figure S.2). Notably, the initial high density of the PBMC culture (∼10^7^𝑐𝑒𝑙𝑙𝑠/𝑚𝑙), mimicking a similar environment as in the lymph nodes, ensures a relatively high viability of CLL cells in both autologous and monocultures (Kim et al., 2024; Wodarz et al., 2014). Indeed, Figure 1 (b) reveals only a slight decay in cell viability in the autologous condition (above 90%), but also high viability for CLL cells in monoculture for two out of three patients, thus in the absence of any other immune cells. Specifically, only patient 2 shows viability below 90%, starting around D8, with one replicate reaching as low as 20%. We, therefore, conclude that the protective effect of immune cells in the culture has a strong impact only on patient 2, for which a clear difference between monoculture and autologous culture can be seen starting from D8, coinciding with previously reported times of formation of NLCs from monocytes initially present in the culture in small percentages.

**Figure 1:**
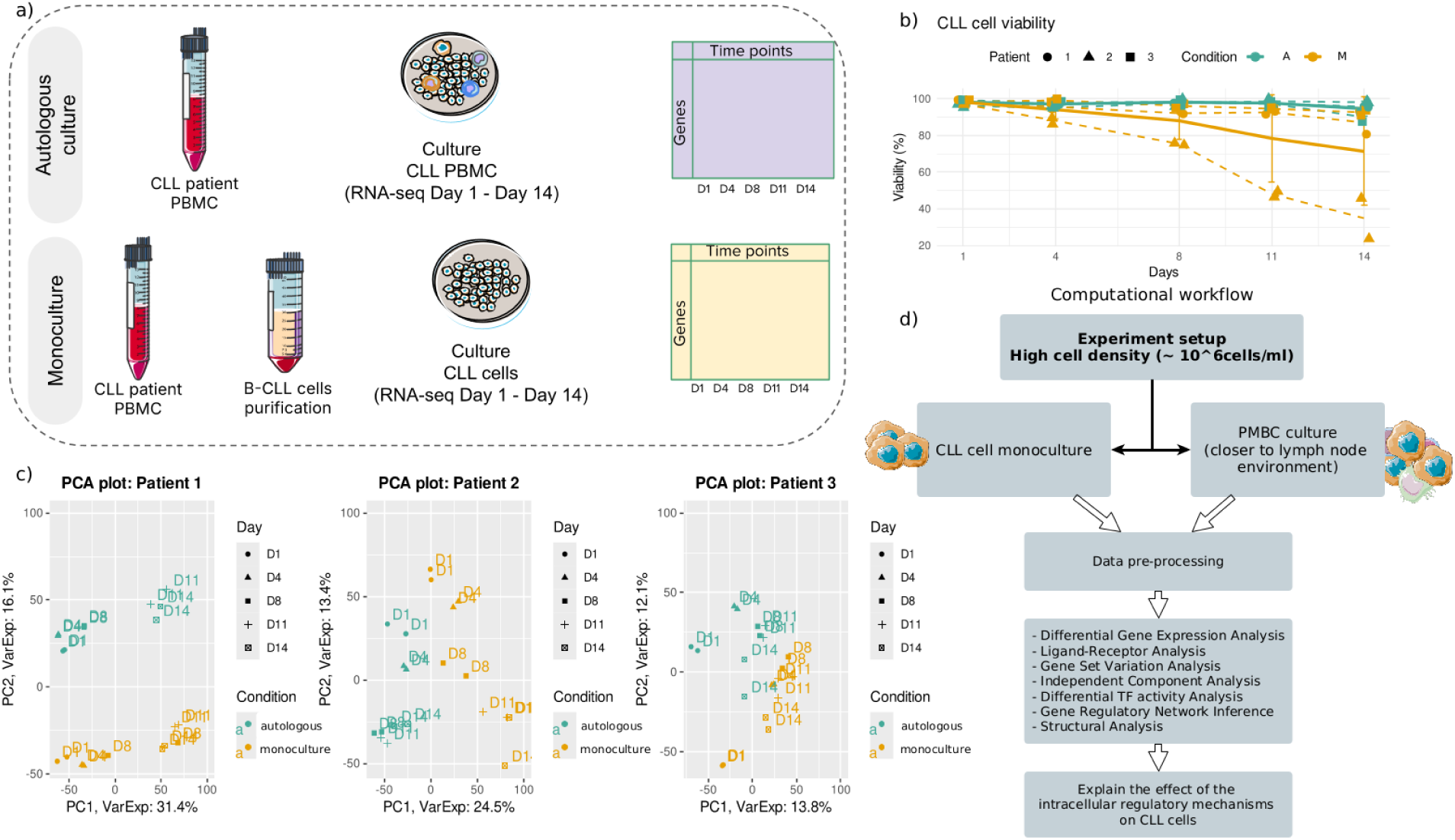
Experimental setup and computation workflow. **(a)** General protocol of *in vitro* experiments of CLL cell cultures in two conditions. **(b)** CLL cells’ viability in autologous culture and monoculture. **(c)** Principal Component Analysis of time-series RNAseq for each patient in monoculture and autologous culture. PC3 is shown in Figure S.3. **(d)** The main computational analyses performed on the time-course RNAseq datasets.

Each generated RNAseq dataset is pre-processed (quality control, normalisation and low variance gene filtering (see <u>Materials and Methods</u>)). We avoid performing any batch correction on the datasets from the different patients to preserve the intrinsic patient variability, allowing us to capture biologically relevant inter-patient differences without eliminating potentially important patient-specific effects. We perform Principal Component Analysis and observe significant differences between patients’ transcriptomes (Figure 1 (c) and Figure S.3). For example, the dominant source of variance could be attributed to the temporal signal in patient 1 (PC1 explaining 31.4% variance), with difference between conditions captured by PC2 (16.1% variance explained), while in patient 2 the difference between autologous and monoculture dominates (PC1; 24.5% variance explained) while PC2 captures temporal differences (13.4% variance explained). Patient 3 shows less clear patterns, with both the first two components explaining less than 14% of variance and both being weakly associated with the culture conditions, while PC3 (explaining 10.9% of variance) tends to separate initial time points from the last two (Figure S.3).

### In-depth analysis of temporal transcriptomic profiles reveals the main differences between autologous and monoculture

We perform an extensive analysis of gene expression profiles of autologous culture and monoculture to quantify the extent to which the CLL cell-intrinsic regulatory mechanisms and survival are impacted by the presence of immune cells in the environment (Figure 1 (d)).

Differential gene expression analysis (DGEA) between the two conditions, as well as between successive time points, is conducted (|𝑙𝑜𝑔_2_𝐹𝐶| > 1. 5, 𝑝𝑣𝑎𝑙 < 0. 05), followed by pathway enrichment analysis on the differentially expressed genes (Figure 2 (a)-(c) and <u>Materials and Methods</u>).

**Figure 2:**
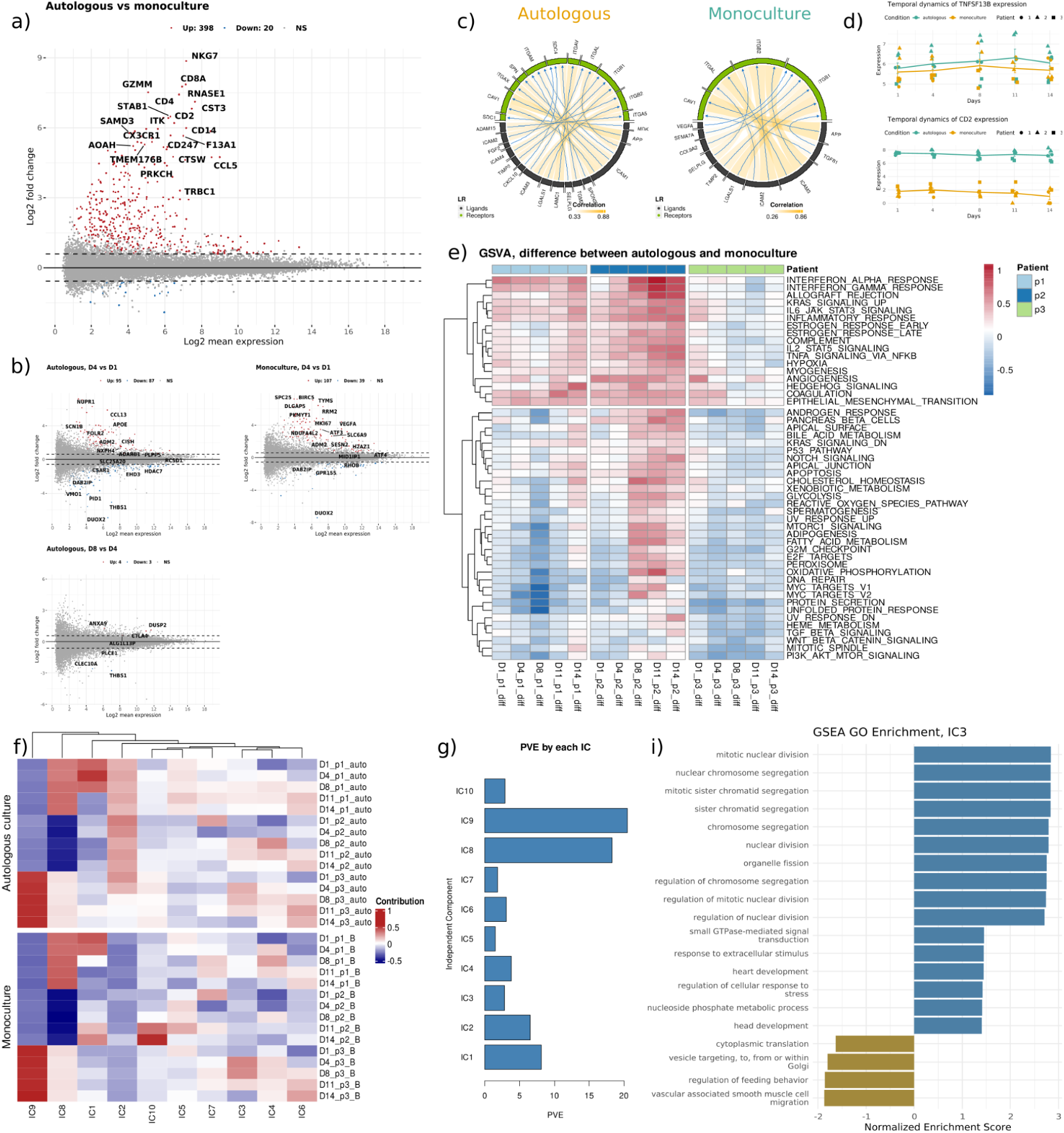
Main results from in-depth computational analysis comparing autologous culture and monoculture. **(a)** Differential gene expression analysis between autologous culture and monoculture. **(b)** Volcano plot highlighting differentially expressed genes between successive time points for autologous culture and monoculture (|𝑙𝑜𝑔_2_𝐹𝐶| > 1. 5, 𝑝*_val_* < 0. 05). **(c)** Ligand-receptor pair analysis of autologous culture and monoculture reveals differences in activated pairs between the two cultures. **(d)** Temporal expression dynamics of TNFSF13B (BAFF) and CD2 ligands in the two culture conditions. **(e)** Difference in Gene Set Variation Analysis results between autologous culture and monoculture. **(f)** Sample scores derived by independent component analysis performed using BIODICA in the two cultures. Different time points in temporal order and patients in rows. **(g)** Barplot showing the percentage of variance in gene expression explained (PVE) by each component. **(i)** Gene Set Enrichment Analysis of IC3 genes.

Amongst the genes most strongly and significantly up-regulated in autologous culture versus monoculture conditions, all time points considered, we identify several genes that are strongly expressed in T cells, NK cells and monocytes (e.g., CD4, CD8A, NKG7, GZMM, CD14) (Figure 2 (a)). Despite the FACS cell proportion and deconvolution estimates suggesting a very low abundance of non-CLL cells in the culture, the results of this differential gene expression analysis could reflect the presence of these cells in the autologous culture samples. This is also reflected in the pathway enrichment analysis of the upregulated genes comparing autologous culture versus monoculture conditions, revealing significant biological processes and pathways associated with the cellular responses of immune cells (Figure S.4 (a)). The most enriched pathways include inflammatory response, chemotaxis, and various aspects of cell motility, such as leukocyte chemotaxis and cell migration, likely highlighting the involvement of immune-related mechanisms, particularly those regulating immune response and cytokine production.

Several signalling pathways are also enriched, including the chemokine signalling pathway, cytokine-cytokine receptor interaction and interleukin signalling, reflecting enhanced immune communication and signalling. These findings collectively are consistent with a robust activation of immune and cellular signalling processes under autologous conditions compared to monoculture (Dubois et al., 2020). In addition, DGEA between the two conditions is performed for each time point separately, revealing that the differentially expressed genes at early time points closely resemble those obtained across the whole time course (complete results can be found in the <u>GitHub repository</u>).

Finally, DGEA between successive time points is performed for each condition separately. In both conditions, we observe the majority of differentially expressed genes at early time point comparisons, mostly related to cell cycle regulation (Figure 2 (b) and Figure S.4 (b), (c)). Considering the expression profiles of the differentially expressed genes across time in the two conditions, we identify a gene module up-regulated only from day 4 and only in autologous culture (cluster 𝑘 = 3, Figure S.4 (d)), mostly enriched in lipid metabolism (Figure S.4 (e)). To further investigate whether the CLL microenvironment contributes to the induction of these genes in CLL cells, we complement our analysis with an external validation using single-cell CITEseq data from PBMC of a CLL patient sample before treatment, under treatment and at relapse (Cadot et al., 2020). We examine the expression of cluster 𝑘 = 3 gene module within this dataset by computing a module expression score across annotated cell populations. We detect a non-negligible expression of the same module in a subset of CLL cells (Figure S.4 (f)), suggesting that, within the autologous culture, the microenvironment may drive CLL cells to induce metabolic reprogramming toward enhanced lipid metabolism, potentially reflecting a more tumour-supportive microenvironment (Rozovski et al., 2016).

To dissect potential signalling interactions between cells in the culture, ligand-receptor (L-R) analysis is conducted, identifying the L-R pairs expressed in the two cultures (Figure 2, (c) and Figure S.5). The results, presented both as L-R pair activation scores or as pathway scores, recapitulate the high patient heterogeneity observed previously. Interestingly, the autologous culture exhibits an expression of ADAM15, FGF2, MDK and CXCL10 ligands, which are not activated in the monoculture, suggesting that cell-cell physical interactions and adhesion, as well as potentially interactions with the matrix, immune crosstalk and activation of BCR signaling, play a crucial role in modulating CLL cell behaviour in the context of the TME (Table 1). Other receptors, such as BAFF (TNFSF13B), associated with CLL-NLC interactions, and CD2, a T cell surface antigen shown to be expressed also by NLCs and interacting with LFA3 (CD58) expressed on CLL cells, mediating the NLC protective effects (Boissard et al., 2016), are also more expressed in autologous compared to monoculture conditions. The difference in BAFF becomes stronger towards the end of the culture, likely revealing CLL-NLC crosstalks (Boissard et al., 2015; Endo et al., 2006; ten Hacken et al., 2014) (Figure 2 (d)) that are activated once the NLCs have formed towards D7-D8 of the culture, corroborating our previous findings (Boissard et al., 2016).

**Table 1:**
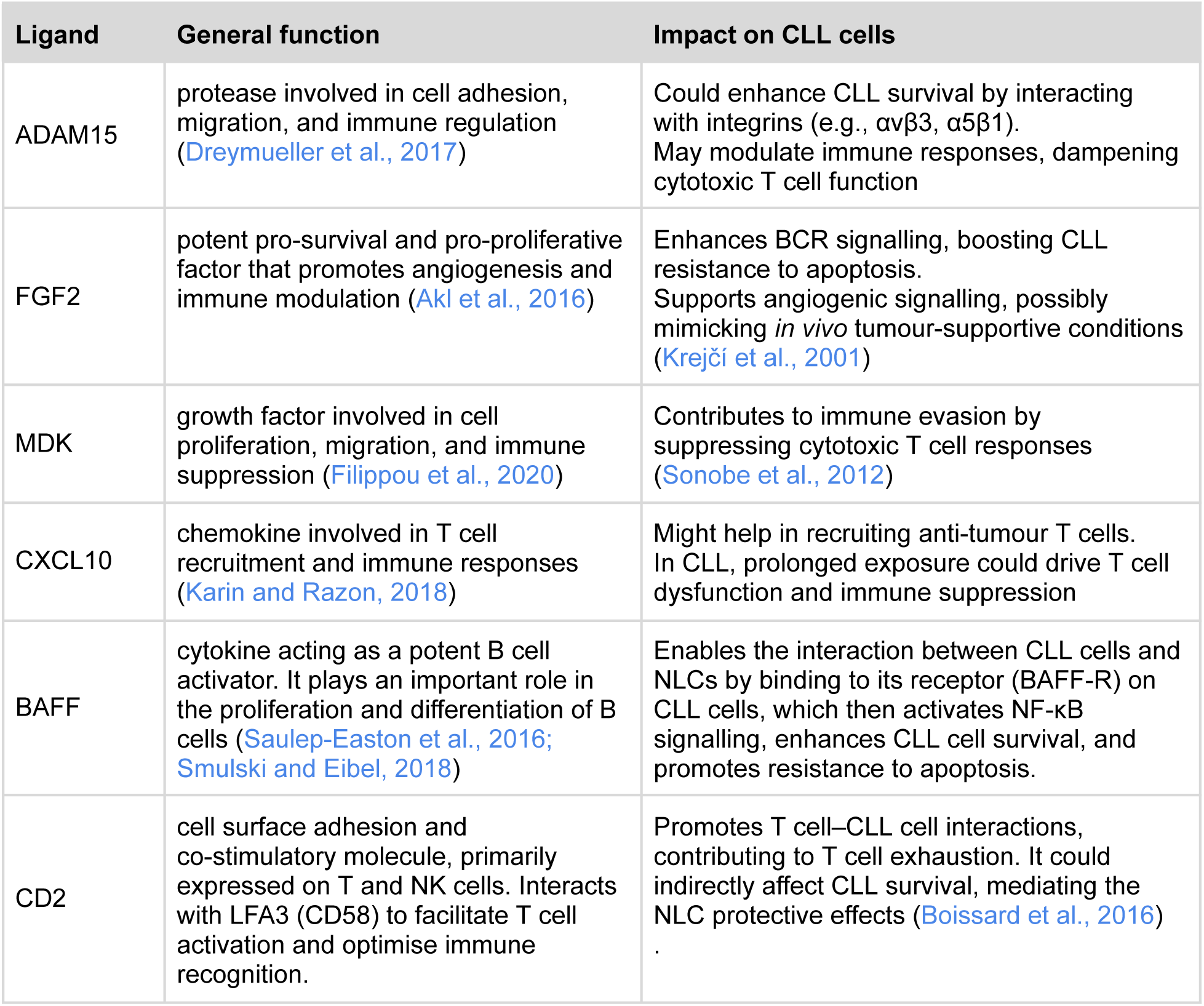
Receptors expressed in autologous culture but not in monoculture and their potential impact on CLL cells.

Finally, Gene Set Variation Analysis using the Hallmarks gene sets extracted from the Human MSigDB Collections is performed to highlight the main differences in pathway enrichment between the two conditions (Figure 2, (e)). Similar to the L-R analysis, the results are impacted by the high patient heterogeneity observed initially. In the autologous condition, immune-related pathways such as inflammatory response, IL6-Jak-STAT3 and IL2-STAT5 signalling, and INFγ response appear consistently enriched, particularly in later time points. This reflects a sustained activation of immune mechanisms over time in the autologous culture compared to monoculture. The variability between patients is evident, with individual patient profiles showing distinct enrichment trends. For example, metabolic pathways such as cholesterol homeostasis, fatty acid metabolism, and glycolysis show late enrichment in patients 1 and 2, whereas in patient 3, their activation is less pronounced or occurs at earlier time points. In addition, patient 2 displays a stronger activation of oxidative phosphorylation or hypoxia-related pathways in autologous culture compared to patients 1 and 3. Pathways related to cytokine production, such as TNFα signalling, also differ significantly between patients and conditions, underscoring the personalised nature of cellular responses to these culture conditions. Exclusively in patient 2, we observe a late differential activation of proliferation pathways (MYC and E2F-related), potentially reflecting the presence of conditions conducive to CLL cell proliferation in the autologous culture.

### Independent Component Analysis decomposes the cellular processes altered in CLL cells in autologous and monoculture conditions

To further dissect the biological processes occurring in the cells in the different conditions throughout the time course, we perform Independent Component Analysis (ICA) using BIODICA (Captier et al., 2022), thus identifying mutually statistically independent sources of variance within the gene expression data (also called Independent Components, ICs). Briefly, ICA performs unsupervised linear separation of mixed signals and quantifies the contribution of individual genes to each component (gene scores or metagenes) and the importance of each component in each sample (sample scores or metasamples). ICA was previously applied to investigate the sources of variability in time-series transcriptomics data (Aynaud et al., 2020). We apply BIODICA to the aggregated dataset, including both autologous and monoculture conditions, to identify ICs that are associated with different patients, conditions and time points.

We compute ICA decomposition into 10 ICs (based on stability analysis, Figure S.6 (a)), as shown in Figure 2 (f) and Figure S.6 (b). In addition, we estimate the proportion of variance associated with each IC in order to quantify the relative strength of each component’s contribution to the overall expression variability (Figure 2 (g)). Two components are clearly associated with patient-specific features (IC8 and IC9), suggesting that the remaining ICs should not be strongly affected by patient heterogeneity and rather capture conditions or time-specific processes. Notably, only IC2 is strongly associated with the difference between the two conditions (autologous versus monoculture), independent of time point, while the remaining components exhibit more complex variation. Considering the pathway enrichment analysis (Gene Set Enrichment Analysis, Reactome database) (Figure S7), revealing processes related to immune activation, and the fact that IC2 sample scores are stable across time, we hypothesise that IC2 is driven by the difference in the composition of the samples (a few non-adherent immune cells other than CLL cells are present in the autologous culture samples). However, the contribution of IC2 to the global transcriptomic variability remains small (Figure 2 (g)), which directs us to focus on those ICs that are related to temporally regulated processes, likely involving the CLL cells’ response to their environment. In particular, IC3 increases specifically in autologous culture in patient 2, peaking between D4 and D8, and is strongly associated with proliferation pathways, as revealed by pathway enrichment analysis (Figure 2 (i)). This leads us to associate IC3 with the pro-proliferation signals coming from NCLs and Tregs in autologous cultures.

Notably, IC10 uniquely characterises patient 2, showing a significant increase in the monoculture, mirroring the decline in cell viability towards the end of the culture for this patient, and likely being associated with the activation of apoptotic pathways. Interestingly, this component captures a negative regulation of IL-10 signalling, BCR receptor signalling, and ion and calcium homeostasis, capturing the reduction in cancer-cell-specific signalling that accompanies the activation of apoptosis. The specific activation of IC10 only in patient 2 reinforces the findings that CLL cells from other patients can indeed survive in the absence of the protective signals present in the autologous culture. This resilience of CLL cells from patients 1 and 3 is likely achieved through autologous signalling from other CLL cells that are found in proximity in such dense culture conditions. Lastly, IC1 also shows a specific increase across time exclusively in the monoculture for patient 2, while it has an almost opposite trend compared to patient 1 in both conditions. The enrichment of IC1 genes suggests involvement in collagen processing and a negative association with apoptosis.

Looking for more global patterns, we notice that IC6 shows an increasing trend in all conditions, patients and replicates and is enriched for processes related to nutrient deprivation, likely reflecting metabolic shifts in CLL cells towards the end of the culture due to reduced nutrient concentration (Figure S7 and Supplementary file).

### Transcription Factor activity estimation identifies differences in signalling pathways between autologous culture and monoculture

Transcription factors (TF) are of particular interest due to their role in the regulation of a number of target genes, but they are rarely regulated at the transcriptional level, often showing very low levels of gene expression. On the contrary, by combining prior knowledge on networks of TFs and their target genes with gene expression data, several algorithms exist to calculate their activity, which can depend on protein levels, post-translational modifications and co-factor interactions. In addition, TF activity calculation is also a way to reduce the high dimensionality of gene expression datasets to around ∼1200 TFs. We, therefore, focus on calculating the activity of all the TFs for each patient in each condition, extracting the most differentially activated (DA) TFs across time points. To identify candidate regulators of the dynamic processes in our cultures, we select only TFs having a significant change between successive time points as DA TFs (Figure 3, (a); more details about the selection can be found in <u>Materials and Methods</u>). In total, a list of 186 TFs is identified as significantly DA.

**Figure 3:**
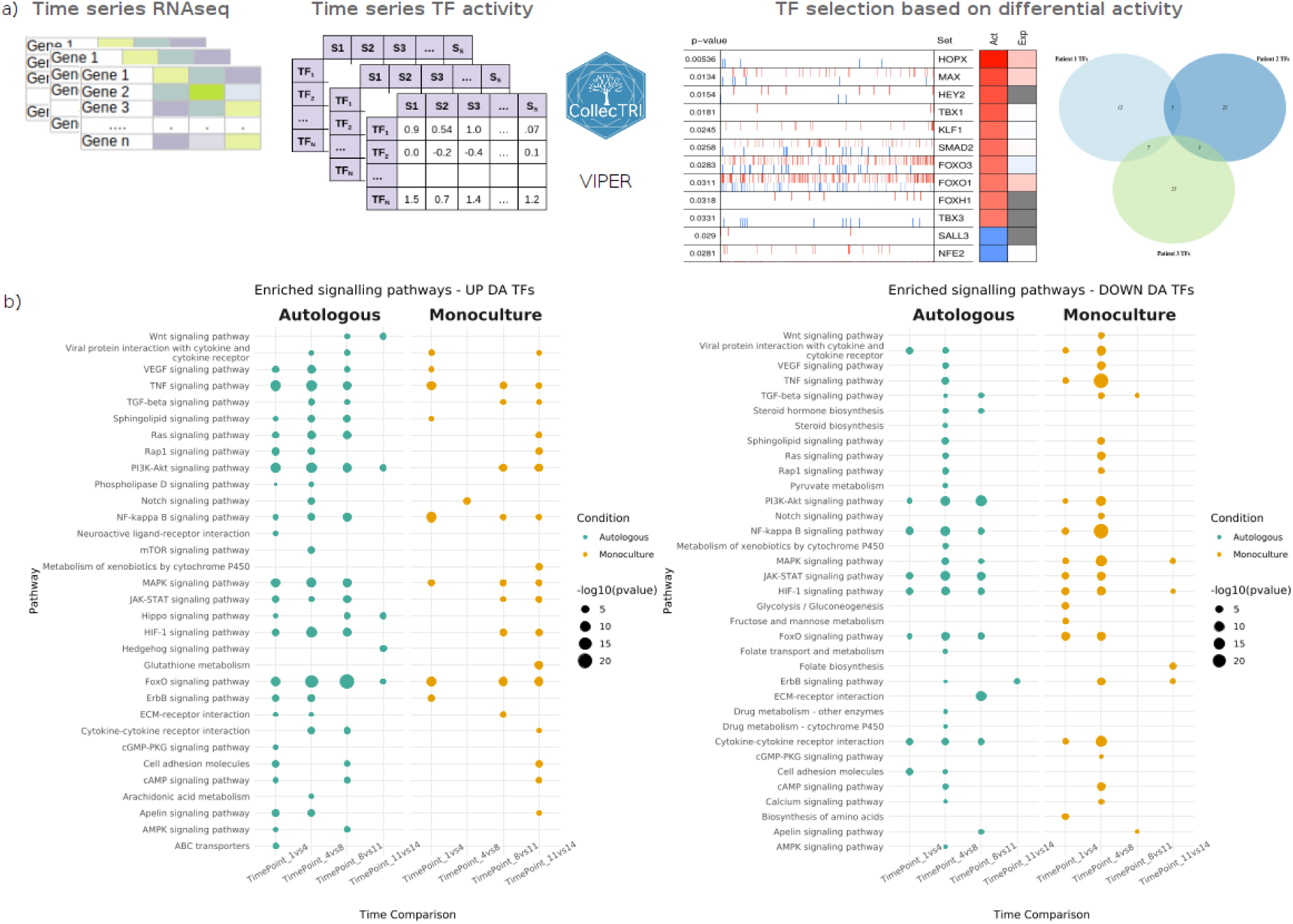
Selection and analysis of the most differentially activated (DA) TFs. **(a)** The selection of the most differentially activated TFs is performed as follows: after calculating the TF activity across time points for each patient and each condition separately (see Materials and Methods), DA TFs are selected by comparing their activity between one time point and the previous one, with a cutoff on the 𝑝*_val_* < 0. 05. The selection is done in a patient-specific and condition-specific context. **(b)** Pathway enrichment analysis of the most DA TFs in autologous culture and monoculture across all time intervals for each patient. The resulting plot is obtained from combining the results for all the patients for pathways involved in signalling.

To further assess the robustness of this TF selection approach, we complement our analysis with an external validation using the single-cell CLL CITEseq data (Cadot et al., 2020). Applying the pySCENIC workflow (Aibar et al., 2017; Kumar et al., 2021) to the CLL cell population of this dataset, we identify a set of master regulators distinguishing pre- and post-treatment CLL sub-populations. Strikingly, we observe a significant overlap (59 TFs, 𝑝*_val_* = 2. 738𝑒 − 04 , hypergeometric test, universe = all TFs) 𝑣𝑎𝑙 between these master regulators and the TFs retained in our study, supporting their central role in CLL cells’ dynamic response to their environment and underscoring the relevance of our selection approach. Full details of this validation are presented in the <u>Supplementary Materia</u>l.

The functional profiles of the TFs and their targets performed by the Over-Representation Analysis (ORA) method highlight distinct transcriptional dynamics between autologous and monoculture conditions, emphasising the influence of the cellular microenvironment on signalling pathways (Figure 3, (b)). Key pathways, including Wnt, NF-kB, and MAPK signalling, are prominently enriched in the autologous condition, emphasising activation of pathways related to tissue homeostasis, inflammation, and stress responses. In contrast, the monoculture condition shows a weaker enrichment profile for up DA TFs, with fewer pathways significantly altered, likely due to the absence of intercellular signalling between cancer cells and immune cells. Shared pathways, such as PI3K-Akt and TGFβ signalling, exhibit differential enrichment patterns, reflecting the distinct regulatory mechanisms operating in each condition. These findings underscore the importance of the microenvironment in modulating transcriptional networks.

### Gene regulatory network inference from bulk gene expression time-courses reveals modular regulatory interactions defining CLL cells’ behaviour

Following the selection of the most differentially activated TFs for autologous culture and monoculture conditions, we seek to infer a GRN, thus unveiling how the regulatory interactions between TFs in CLL cells define their phenotypes. To do so, the TF-TF networks for each biological condition are inferred using the algorithm proposed in the dynGENIE3 method (Huynh-Thu and Geurts, 2018), but using as input the TF activity estimates instead of the gene expression values. Details on the method’s algorithm and parameters are provided in the <u>Materials and Methods</u>. The algorithm generates an output file listing regulatory genes, target genes, and their associated interaction weights (𝑤_𝑖𝑗_ ∈ [0, 1]). To select the most significant interactions, we applied filtering of the TF-TF interactions, considering only those with an inferred interaction weight 𝑤_𝑖𝑗_ ≥ 0. 01. Aiming to build a general network that would capture key regulatory mechanisms of CLL cells in the two conditions, condition-specific TF–TF networks are then merged, yielding a network of 186 nodes and 4280 edges. In parallel, to capture the functional relationships between TFs that are differentially active across the same temporal intervals, we consider differential activity scores of each TF between two consecutive time points in a specific condition. We then calculate the correlation of these scores for all TF pairs, keeping only those with absolute correlation |𝑐𝑜𝑟_ij_ | ≥ 0. 5 (see <u>Materials and Methods</u>), from which the interaction signs (positive ≡ activation, or negative ≡ inhibition) are also extracted. This procedure generated a second set of condition-specific networks, which includes 182 nodes and 2614 edges. Finally, we consider the overlap of these two independently derived TF-TF networks, resulting in a consensus network comprising 167 nodes and 498 edges. This hybrid approach captures a spectrum of regulatory interactions by integrating both non-linear (from the dynGENIE3 algorithm) and linear (from the temporal correlation analysis) time-aware relationships.

The inferred network represents context-specific regulatory interactions that shape the response of CLL cells to their *in vitro* environment and, as such, cannot be directly compared to existing networks for B cells or cancer cells that more often capture oncogenic transformation or other types of dynamic phenotype shifts. We, therefore, proceed to validate the network by investigating the biological relevance of its characteristics. Specifically, mapping the information about the TFs on the TF-TF network, namely in which time interval and in which patient the TFs change their activity, or in which condition the TF is selected as variable, reveals interesting patterns in its structure and connectivity (Figure 4 (a), Figure S.8 (a)). Firstly, we observe a segregation of TFs selected in the different culture conditions in distinct regions of the network, where TFs common to both conditions are found to be more centrally located. Interestingly, we also observe a pattern in the centrality of TFs annotated by having differential activity at early or late time points, as evidenced by decreasing betweenness centrality for TFs activated later in the time points, with the TFs that show differential activity in multiple time intervals being the most central of all. Biologically, betweenness centrality reflects the potential of a TF to act as a key regulatory intermediary. This suggests that initial stages in the dynamics of the culture are captured by a core set of TFs, which are also common to the two conditions, while the effect specific to the presence of immune cells in the autologous culture becomes stronger with time. Moreover, the TFs common to several patients are found to be intermediate nodes connecting groups of patient-specific TFs, as evidenced by their betweenness centrality (Figure 4 (b)). This suggests that the processes activated in different patient samples are captured by distinct regions of the network.

**Figure 4:**
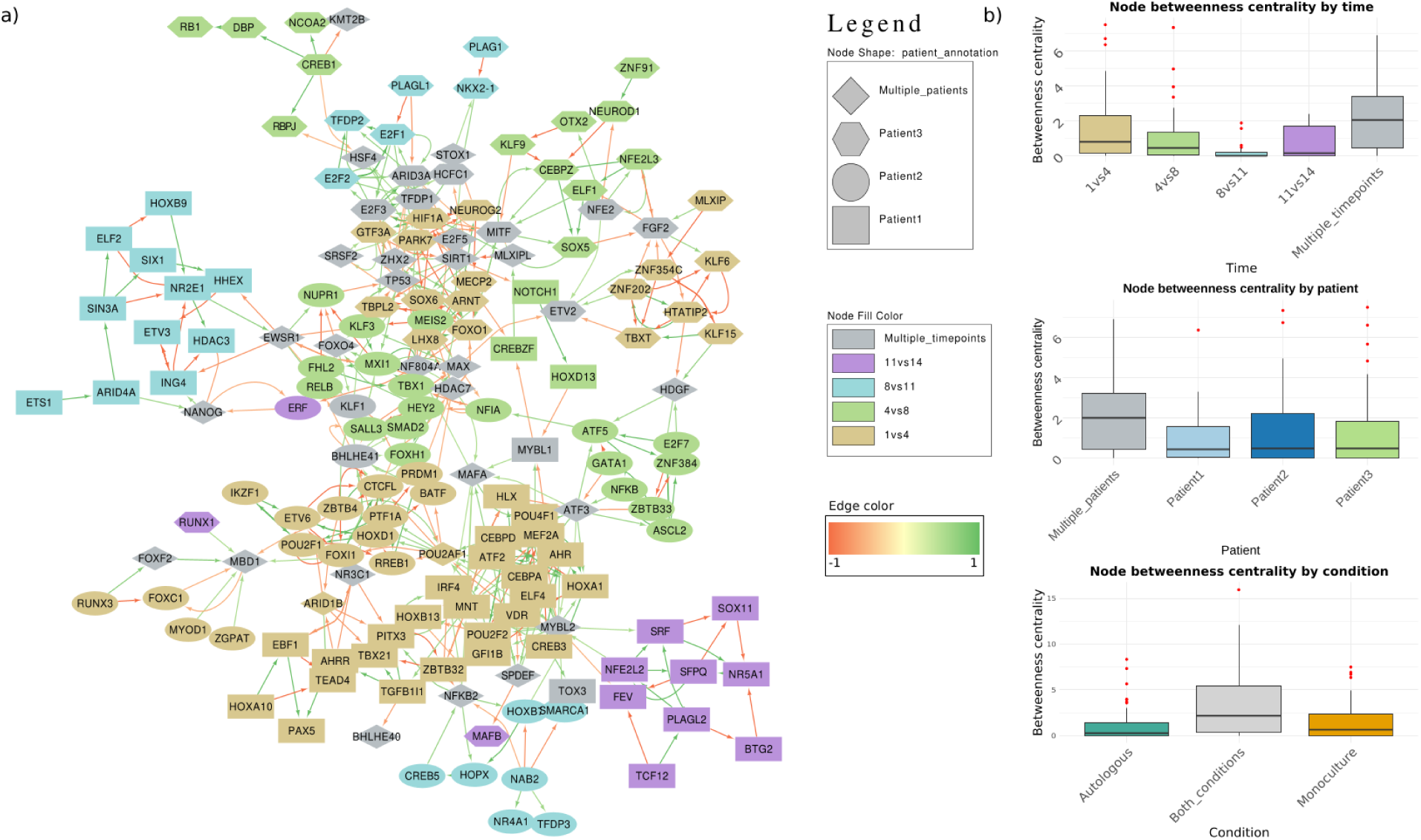
The inferred TF-TF regulatory network and structural analysis. **(a)** Inferred regulatory network of TF-TF interactions. The node features include the time interval in which the TF had differential activity and the patient in which the TF activity specifically changed, whereas the edges’ weight and colour represent the correlation of activity of the two adjacent TFs. **(b)** Betweenness centrality of nodes according to the TFs features (condition, time and patient).

Interestingly, the subset of TFs overlapping with those identified by pySCENIC in the Cadot et al. dataset is found to localise preferentially in the core of the inferred GRN (<u>Supplementary Material</u> and Figure S.9). These TFs are enriched among those activated at early time points, suggesting that the difference between CLL cells before and after treatment could be similar to the changes we observe in our *in vitro* cultures in the first few days.

The network structure shows a tendency of the nodes to be connected to other nodes that are similar with respect to a given feature. To quantify this, we estimate the network assortativity on node features, defined as the correlation 𝑟 (𝑟 ∈ [− 1, 1]) of node features (Table 2) and compare the assortativity from 100 random networks generated from the TF-TF network by shuffling the node labels, thus preserving the topology and the features of the network. Assortativity, or homophily, measures the tendency of nodes in a network to connect with other nodes with similar features. A positive assortativity indicates that nodes with similar features are more connected together, while a negative assortativity implies that nodes with a given feature tend to interact with nodes with other features (Newman, 2003). Our results show a significantly higher assortativity, indicating preferential interactions of TFs that are specific to the same patient, time point or condition. The existence of patient-specific neighbourhoods in this network is potentially linked to the specific transcriptional and regulatory mechanisms and processes activated in each patient. In addition, this might reflect differences in immune responses, metabolic states, or other system-level variations between patients that are intrinsic to the CLL cells. In the context of CLL, the high assortativity may suggest that the phenotypic changes in CLL cells follow unique trajectories in each patient.

**Table 2:**
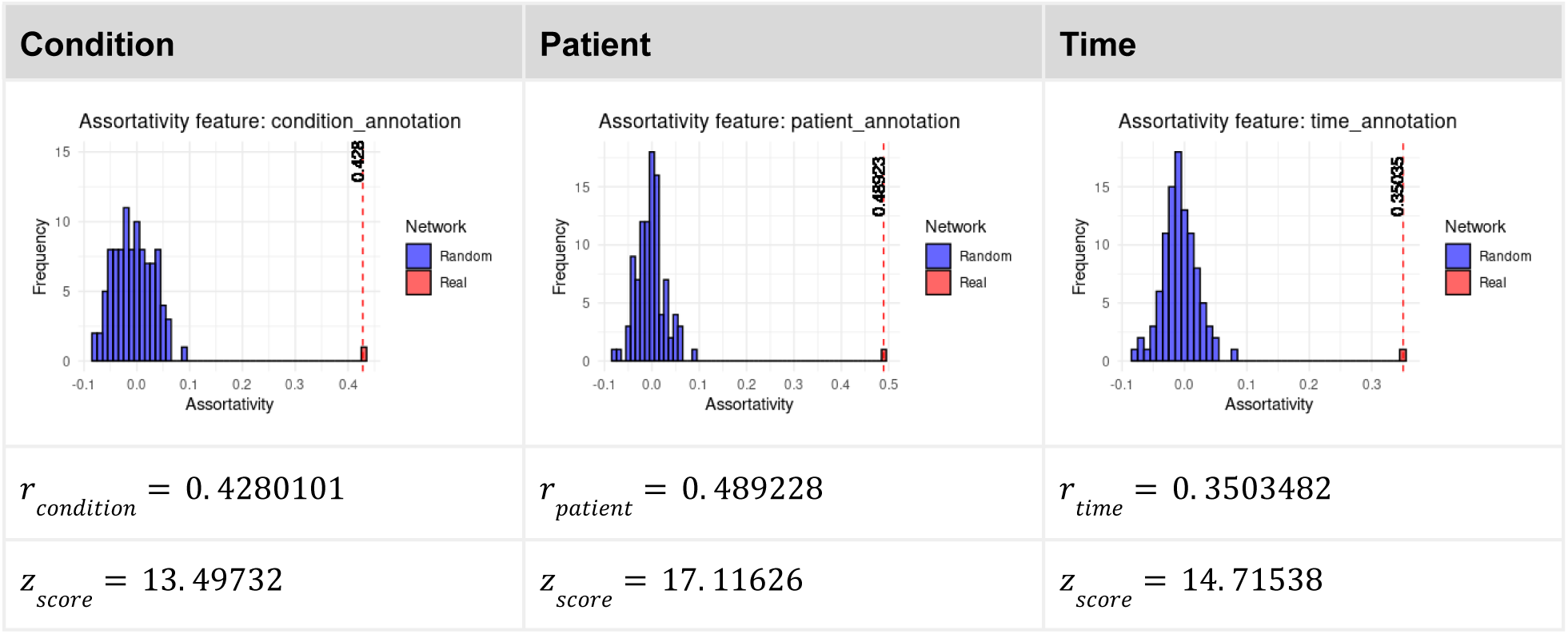
TF-TF network assortativity compared to random network.

To assess whether functionally related TFs participating in specific biological processes form modules in the TF regulatory network, we perform community detection and investigate the modular structure of the TF-TF network (Figure S.8 (b) and <u>Supplementary Material</u>). Comparing the modules with other features of the network, we observe that, apart from the two biggest modules (module 2 and 10) having mixed features, the rest of them capture mostly patient-specific features at different time points and different conditions (Figure S.10), suggesting that the modules represent specific processes taking place in different patients, time points and conditions. For this reason, we performed functional enrichment analysis on each module, expanded with the targets of the TFs (Figure S.11). We observe that module 2 is enriched for pathways involved in immune response, inflammation, and metabolic regulation, including cytokine signalling, interleukin pathways, leukocyte activation, and lipid metabolism. The presence of pathways such as PI3K-AKT signalling, TNF signalling, AGE-RAGE signalling, and IL-10 signalling suggests that this module may reflect the activation of mechanisms in CLL cells related to their survival, immune modulation toward pro-tumoural phenotypes, and resistance to apoptosis, as well as metabolic shifts. On the other hand, module 10 is significantly enriched in pathways related to cell cycle regulation and TP53 signalling, potentially related to cell fate decisions, tumour suppression, and DNA damage responses (Figure S.11).

Looking for more feature-specific modules, we investigate the rest of the modules, classifying them by patient. Interestingly, we observe that each module represents specific processes taking place at different time points and conditions. For patient 1, module 3 consists of TFs from early time points (D1-D4) in the autologous culture and is enriched in B cell activation, differentiation, and cytokine signalling pathways, including JAK-STAT and IL-4/IL-13 signalling, suggesting CLL cells-immune cells interactions. Module 11, consisting of TFs from D8-D11 in the same condition, is primarily enriched in cytokine signalling, PI3K-AKT pathway, and activation of proliferation pathways in a supportive environment. In contrast, module 1, consisting of TFs from D11-D14 in monoculture, is associated with metabolism and stress response pathways, highlighting a shift toward stress adaptation and metabolic reprogramming in the absence of immune pro-tumoural stimuli and nutrient deprivation.

For patient 2, module 4 consists of TFs from two early intervals (D1-D4 and D4-D8) and predominant in monoculture, with enrichment of differentiation and cytokine signalling pathways, reflecting intrinsic programs in CLL cells, independent of autologous immune cell interactions. Module 5, consisting of TFs from D4-D8 in autologous conditions, includes negative regulation of programmed cell death and apoptosis, as well as IL-4/IL-13 and FOXO signalling pathways, indicating a survival-oriented program. Additionally, module 6, consisting of TFs from the beginning of the autologous culture, is enriched in proliferation, differentiation, and cytokine signalling pathways, indicating early activation of proliferative and differentiation programs in CLL cells in a supportive environment.

For patient 3, module 7, consisting of TFs from D1-D4 in autologous culture, reflects activation of proliferation, stress response and metabolic pathways, likely associated with early CLL cell adaptation to the *in vitro* environment. Likewise, module 9 in monoculture shows enrichment of similar pathways. Module 8, in monoculture during D4-D8, is enriched in metabolism, HIF1α, and FOXO signalling pathways, and indicates metabolic reprogramming and stress responses in CLL cells.

## Conclusions and discussions

In this study, we leverage time-series transcriptomics and data-driven GRN inference to investigate how the dynamic interactions between CLL cells and surrounding immune cells determine CLL survival and responses in an *in vitro* TME model. We were initially surprised to notice that only one out of the three patient samples examined (patient 2) displayed reduced survival of isolated B cells compared to B cells cultured within the corresponding PBMC. Indeed, for the remaining two patients, the CLL viability is stable at above 90% across the entire time course, suggesting that autologous interactions between CLL cells cultured at high densities can be sufficient to ensure their survival in some patients.

Our analysis reveals significant differences between monoculture and autologous culture of CLL cells, highlighting the impact of immune cell presence on CLL cell phenotype, gene expression dynamics, and regulatory interactions. Specifically, the differential gene expression analysis demonstrates that autologous culture exhibits upregulation of pathways associated with immune response, cytokine signalling, and metabolic shifts, suggesting a distinct transcriptional landscape compared to monoculture conditions. Similar results are also obtained from the ligand-receptor interaction analysis, which uncovers specific signalling interactions present in autologous culture, with ligands such as ADAM15, FGF2, MDK, and CXCL10 being specifically activated, pointing to immune cell-mediated processes in CLL elicited by the TME. Additionally, a specific upregulation of the BAFF and CD2 ligands in the autologous culture suggests a potential response of CLL cells to the NLC.

Patient heterogeneity is a key determinant feature, clearly underlying heterogeneous CLL cell behaviour, as shown in most results (GSVA, ICA, LR analysis), revealing patient-specific differences in immune activation, metabolic adaptation, and stress response pathways.

While differential gene expression analysis across autologous and monoculture reveals a potential confounding effect on the immune activation signals identified specifically in autologous culture of the presence of a few non-CLL cells in the samples analysed (estimated by FACS and a deconvolution approach to be <10%), we employ ICA to distinguish the different processes that underlie the transcriptomics profiles we record for different patients.

ICA analysis identified one component strongly associated with the patient sample (IC9), one separating patient 3 from the other two (IC8, which might be related to the fact that the patient 3 sample is the only one that had not undergone freezing), and one component that distinguishes autologous culture and monoculture without strong changes across the time course (IC2). As expected, the strongest dynamic differences between monoculture and autologous culture, attributable to the protective effect of specific immune cells in the culture, are mostly seen for patient 2. In particular, IC3, which is related to cell proliferation, seems to be increasing specifically in the autologous culture around day 8, consistent with an effect of NLCs having formed and providing pro-tumoural signals in combination with the presence of Treg cells that can enable cell proliferation *in vitro*. This effect is potentially mediated by the BAFF ligand, which is seen to increase specifically at this time point. On the contrary, IC10 and IC1 display a specific upregulation in the monoculture of patient 2, suggesting the activation of apoptosis and collagen remodelling purely in the absence of supportive immune cells. Finally, IC6 displays a global upward trend across the time course in all patients and conditions, the strongest upregulation being for the autologous condition of patient 2, apparently related to starvation as a consequence of lack of nutrients at the end of the culture. Presumably, the cells of patient 2 in autologous culture might be more acutely affected by this stress due to their increased nutrient demands as a consequence of the activation of proliferation pathways.

Furthermore, GRN inference uncovers modular regulatory interactions in a TF-TF network defining CLL behaviour. We observe strong assortativity of nodes’ features describing the conditions, time points and patients in which the TFs change their activity, reinforcing the idea that the topology of the network can capture the effect of the environment, the timing of different processes and inter-patient heterogeneity. These differences can be understood when considering the underlying interpatient genetic and non-genetic heterogeneity that is typical of cancer, combined with potentially heterogeneous composition and characteristics of the PBMC in each patient. In addition, these findings emphasise the importance of considering the TME in studying CLL dynamics, particularly at the cellular level, where the characteristics of the surrounding environment (e.g. density of the cells, presence of other immune cells, hypoxia, lack of nutrients) significantly impact cancer cell survival.

The study also highlights the potential of data-driven GRN inference approaches in identifying novel regulatory interactions that could help to identify new therapeutic targets. The relevance of the network we have inferred is reinforced by external validation using pySCENIC on a single-cell RNAseq dataset from PBMC of a CLL patient, which we use to identify master regulators in these cells (<u>Supplementary</u> <u>Material</u>). We identify a conserved subset of regulators that overlap between master regulators in the single-cell RNAseq data and our TFs, inferred to be key for the response of CLL cells to immune cells in the *in vitro* culture. These TFs occupy topologically central positions within the network, consistent with regulators orchestrating early transcriptional programs in our culture system. Furthermore, we detect an upregulation of genes induced in the culture of treated CLL cells in the study by Cadot et al., raising the intriguing possibility that therapeutic perturbation may elicit transcriptional effects akin to those observed in our *in vitro* culture of PBMCs.

Lastly, computationally, the ensemble of analyses presented constitutes a workflow to go from time-series RNAseq datasets to GRN inference, providing a detailed investigation of the data at different levels. The methodological framework established in this study, from data analysis to network inference, contributes significantly to the broader field of GRN inference by providing a scalable and adaptable workflow for studying gene regulation in time-resolved contexts. This framework is particularly relevant for diseases characterised by dynamic changes in cell states, where regulatory networks are continually reshaped by both intrinsic cellular processes and extrinsic environmental factors. In particular, the selection of the most differentially activated TFs and the combination of gene expression with the TF activity for the network inference provide an unbiased approach to data-driven network inference while integrating knowledge from different sources.

### Limitations of the study

Despite the value of this data-driven approach to describe CLL dynamics in different environments, there are some important limitations in this study that we will list below.

First, as far as the experiments are concerned, these *in vitro* cultures might not fully represent what is happening in the patient lymph nodes, where the density of CLL cells is as high as what we reproduce in the cultures, but potentially different cell populations, particularly immune cells, can be present in different proportions and within a different physical environment. For example, in our cultures, monocytes differentiate into macrophages in contact with the culture plate, something which is clearly artificial and potentially different from the differentiation of monocytes upon entering a tissue.

Second, heterogeneity across patients is well known in these cultures, the polarisation of macrophages into NLCs being extremely patient-specific in both time scales and phenotypes of NLCs produced (Domagala et al., 2022; Verstraete et al., 2023). This heterogeneity could strongly affect the generalisability of our results.

Third, it is worth noting that during the experiment, there has not been any addition of nutrients in the *in vitro* culture. Therefore, toward the end of the experiment, the cells might be facing starvation due to a decrease in the overall nutritional level, with the activation of stress response pathways. However, we avoid replenishing the culture media in order to prevent introducing a perturbation at regular intervals across the time course that would perturb the CLL cells’ dynamics, including a dilution of the concentrations of cytokines produced by the cancer cells.

Fourth, and most importantly, the presence of non-CLL in the culture, even in low percentages, can introduce alterations in gene expression profiles that are caused by phenotypic switches in non-CLL cells, which could be confounded with the CLL cells’ transcriptional dynamics. Such contamination risks could raise concerns about whether the inferred regulatory networks capture CLL-intrinsic responses to microenvironmental cues or whether they reflect processes occurring in non-CLL cells that are present in small proportions in the sample. To address this, we quantify the strength of this potential confounding effect in two ways: first, we perform gene expression deconvolution of our bulk RNA-seq data using CLL-specific signatures derived from the single-cell dataset of Cadot et al; second, we investigate whether the genes that we observe to be activated in our time course specifically in the autologous condition, are also found expressed in the CLL cells extracted from this single-cell RNAseq dataset. Indeed, these genes, involved in lipid metabolism, are expressed in subsets of CLL cells from Cadot et al (Figure S.4 (f)). Careful consideration of these genes highlights the presence of FOLR2 and APOE, which are genes often associated with myeloid cells (Domagala et al., 2025; Guimarães et al., 2024; Tao et al., 2024). Prior reports describing the upregulation of T cell and myeloid-specific transcripts within purified or clonally tracked CLL cells (Dong et al., 2020) reinforce our interpretation that the phenotype alterations in CLL cells that we capture based on bulk transcriptomics are *bona fide* CLL processes rather than artefacts of contamination.

To our knowledge, no previous study has monitored the transcriptome of CLL cells cultured with cognate PBMC for close to two weeks. Under these conditions, we observe pronounced CLL phenotypic plasticity, consistent with prior reports describing that CLL cells can adopt transcriptional programs of other lineages. For example, T-BET (TBX21), classically a master regulator in T cells, can be induced in malignant B cells, where it exerts tumour-suppressive functions (Roessner et al., 2024). Similarly, CLL cells have been shown to activate myeloid programmes (Dong et al., 2020), and this might explain the emergence of secondary myeloid malignancies in some patients (Bigenwald et al., 2025). Similar lineage flexibility has been described in other B-cell malignancies, such as clonally related follicular lymphomas and histiocytic/dendritic cell sarcomas (Feldman et al., 2008). At the mechanistic level, disruption of the B-cell TF PAX5 or enforced expression of C/EBP TFs can reprogram mature B cells toward a myeloid fate (Cobaleda et al., 2007; Rapino et al., 2013), underscoring that the atypical transcriptional programs we observe likely reflect intrinsic plasticity of CLL cells rather than technical artefacts. Further investigation using single-cell studies on similar *in vitro* co-cultures will be key to understanding the causes of a high expression of these genes in CLL cells, but our investigation of single CLL cells from Cadot et al. demonstrates that these genes can be expressed by CLL cells.

Finally, the use of samples from three distinct patients allows us to uncover patient-specific regulatory patterns and highlight the context-specific nature of CLL cell behaviour. However, the small sample size limits generalisation, and caution is warranted in drawing broad conclusions. Future studies involving larger patient cohorts or employing more stringent cell sorting or single-cell approaches will be essential to confirm the reproducibility of these findings and to disentangle patient-specific effects from more generalizable regulatory mechanisms.

Regarding the GRN inference, clearly, the choice of the GRN inference method directly impacts the results regarding the inferred TF-target interactions. Data-driven GRN inference remains an open challenge in computational biology, with limitations coming from various factors, such as incomplete multi-omics datasets, a high fraction of false positive interactions, difficulty in merging samples from different patients due to patient variability, etc. (Henao et al., 2022; Kernfeld et al., 2023; Marku and Pancaldi, 2023). In our case, the algorithm is run for each condition separately but considering all the patients to capture global non-patient-specific processes across time. While GRN inference algorithms employ gene expression profiles, we are interested in capturing a network of regulators more easily identified by TF activity analysis. We thus combine the dynGENIE3 algorithm with a more *ad hoc* approach, capturing the correlated changes in TF activities across successive time points. Furthermore, as the inference method gives the highest performance when it integrates several datasets compared to the scores obtained when only one dataset is exploited (Marbach et al., 2012), it suggests that datasets from different patients containing different and complementary information should be exploited jointly. To further assess the biological relevance of the inferred regulatory interactions, we compare our set of TFs with those identified by pySCENIC in an independent CLL single-cell dataset (Cadot et al., 2020), observing a substantial overlap, particularly among TFs central to our network and active at early time points (see <u>Supplementary Material</u>). Lastly, validating the inferred network would include comparison with a gold-standard network or performing additional experiments to study the effect of perturbations of nodes or their downstream targets (Fleck et al., 2023; van der Wijst et al., 2018). In our case, no clearly matching gold-standard GRN exists, limiting our options for the validation procedure. We thus aim to ensure the biological plausibility of the inferred interactions, relying on network topology and enrichment of known signalling pathways.

In the future, experimental perturbation studies, such as drug treatments, will be critical for validating predicted causal links. Such perturbation-based validations, combined with single-cell techniques like CITE-seq or ATAC-seq to track functional consequences on chromatin accessibility and protein expression, could provide orthogonal support for the GRN structure. In addition, by combining the data-driven and prior knowledge approaches through incorporating curated regulatory interactions from databases, such as TRRUST (Han et al., 2018) or collecTRI (Müller-Dott et al., 2023), we can enhance network accuracy and identify conserved regulatory mechanisms across different biological conditions. Future efforts should focus on integrating computational and experimental validation strategies to build robust and biologically meaningful GRNs that can provide actionable insights into CLL progression and therapeutic targeting.

### Perspectives

Building on the inferred gene regulatory network, this study opens the way for applying dynamical modelling approaches to simulate the temporal behaviour of CLL cells. In particular, Boolean modelling (Hall and Niarakis, 2021; Marku et al., 2023) offers a powerful framework to explore how internal and external signals propagate through the network to drive transitions in cellular phenotypes. Similarly to using expression data for GRN inference, the information in the changes of the genes’ expression is also used to infer the Boolean rules that govern these changes (Barman and Kwon, 2018; Gjerga et al., 2020; Hall and Niarakis, 2021; Henao et al., 2022; Hérault et al., 2023; Ostrowski et al., 2016; Paulevé et al., 2020; Razzaq et al., 2018) and simulate network behaviour under various perturbations (Naldi et al., 2018). Such models can help identify critical nodes or combinations of regulators that shift the system toward specific cell states. Future integration of this data-driven GRN with Boolean modelling could thus provide mechanistic insights into how CLL cells adapt to different microenvironmental contexts and suggest candidate targets for therapeutic intervention.

## Materials and Methods

### Experimental design

Two different types of cell cultures are used for this study. **Autologous culture**: This culture comprises all peripheral blood mononuclear cells (PBMCs). The culture generation protocol for this category involved the collection of samples from three separate patients. Additional clinical information about the patients’ age, sex, mutation, treatment and copy-number alterations can be found in the Supplementary material, Table S1. Each patient’s samples underwent biological duplication, generating two distinct replicates per patient. **B-CLL monoculture**: B-CLL cells are isolated and cultured individually.

PBMCs are isolated from the blood of CLL patients by Ficoll gradient (Ficoll-Paque Plus™, Thermo Fisher Scientific, Illkirch, France). To generate the autologous culture, the B-CLL cells are cultured at 10^7^ cells/mL in RPMI 1640 GlutaMax™ supplemented with 10% Fetal Calf Serum and 1% Penicillin/Streptomycin (Gibco/Life Technologies, Courtaboeuf, France) in a 6-well plate for 14 days. The culture media is not replaced to avoid introducing perturbations in the natural dynamics of the culture. At 5 different time points (D1, D4, D8, D11 and D14), the cells in suspension are collected and preserved as a dry pellet, which contains CLL cells and a small number of T cells, NK cells and potentially traces of monocytes at the first time point, before they have differentiated into macrophages that are adherent to the plate. In parallel, we did a control condition where we only had the isolated B-CLL cells. To do so, we isolate the B-CLL cells by enrichment using a magnetic kit (EasySep™ Human B Cell Enrichment Kit II Without CD43 Depletion, Stemcell, Saint-Egrève, France) and culture them in the same conditions as the autologous culture. When all the time points are collected, we proceed to the extraction of the RNA from the dry pellets generated for each, using the RNA extraction kit (RNeasy Mini Kit, Qiagen, Courtaboeuf, France). RNA sample qualification and sequencing are performed using the same protocol as in autologous cultures.

### Data processing: quality control, data normalisation and low-count gene removal

Genes with little or no expression information (e.g., rows with zeros or constant expression across the time series) are removed in a first filtering step, and only genes with a count of 2 or more in at least two samples are retained. Normalisation is performed using DESeq2 (Love et al., 2014). Differential expression analysis between conditions or between successive time points in specific conditions is also performed with DEseq2 (|𝑙𝑜𝑔_2_𝐹𝐶| > 1. 5, 𝑝𝑣𝑎𝑙 < 0. 05).

### Ligand-receptor interaction analysis

BulkSignalR (Villemin et al., 2023) is employed to infer ligand-receptor interactions (LRIs) from bulk transcriptomic data by integrating ligand and receptor expression levels with downstream pathway activities. The method identifies significant ligand-receptor-pathway (L-R-pw) triples where the ligand (L), receptor (R), and associated pathway (pw) exhibit correlated activities across the dataset. Potential LRIs are sourced from the *LRdb* database, and pathways are derived from Reactome and Gene Ontology biological processes. The significance of these triples is assessed using a statistical model that considers the null distributions of L-R and R-target gene correlations, pathway sizes, and the total number of target genes. This approach allows for the inference of LRIs without prior knowledge of sample clustering, making it suitable for analysing datasets with unknown or complex cellular compositions.

In our case, using the expression data from both autologous and monoculture conditions, we perform a comparative analysis of LR scores or pathway scores for each sample under each condition. To facilitate the visualisation of the results, we calculate the differences between autologous culture and monoculture conditions and select the most variable LR pairs or pathways.

### Independent Component Analysis (ICA)

We perform ICA with BIODICA (Captier et al., 2022) on the aggregated dataset of autologous and monoculture conditions to identify independent components (ICs) that represent statistically independent sources of transcriptomic variability, elucidating the primary differences between cellular phenotypes in the two experimental conditions. Briefly, each IC is characterised by component loadings, indicating the contribution of individual genes to each component (gene scores) and the association of samples to each component (sample scores). The output includes a metagene matrix, which quantifies the contribution of each gene to the ICs, and a meta-sample matrix, which contains the expression profiles of the ICs across the different cultures, patients and time points.

To quantify the biological relevance of each IC, we compute the proportion of variance explained (PVE) in gene expression and rank genes by absolute loading values per IC. The PVE is estimated by projecting the gene expression data onto the ICs and computing the variance captured. Specifically, given a gene expression matrix 𝑋̂ (genes × samples) and a metagene matrix 𝑀 (genes × ICs, containing loadings), the projection is performed using the Moore-Penrose pseudoinverse:

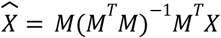

where 𝑋̂ represents the reconstructed expression matrix based on a given IC. The variance explained by an IC is computed as the sum of variances 𝑋̂ across genes, normalised by the total variance in 𝑋̂:

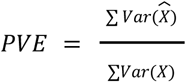

### Transcription factor activity estimation

Virtual Inference of Protein-activity by Enriched Regulon (VIPER) analysis (Alvarez et al., 2016) is used to infer transcription factor (TF) activity across the three experimental conditions, utilising the collection of transcriptional regulatory interactions (CollecTRI) (Müller-Dott et al., 2023) from OmniPath, accessed through the decoupleR package (Badia-i-Mompel et al., 2022). Differential TF activity analysis is performed using msVIPER, a variant of VIPER that evaluates TF activity changes based on a differential expression-derived signature (Alvarez et al., 2016).

To construct this differential signature, we apply a row-wise *t-test* comparing gene expression between conditions. The resulting p-values are converted into z-scores, which serve as input for msVIPER. Additionally, a null model is constructed using 1000 permutations of the dataset to estimate the background distribution of TF activity scores. Differentially activated TFs are identified based on msVIPER enrichment scores, applying a significance threshold of 𝑝𝑣𝑎𝑙 < 0. 05. The Over-Representation Analysis is done using the DA TFs and their target genes from the OmniPath network. Finally, the list of TFs used to generate the gene regulatory network (GRN) is compiled by merging the differentially activated TFs from each patient across both conditions.

### Gene regulatory network inference from TF activities with dynGENIE3

We select dynGENIE3 (Huynh-Thu and Geurts, 2018) for GRN inference due to its strong performance on time-series gene expression data, consistently ranking among the top methods in benchmarking studies such as the DREAM challenges (Marbach et al, 2012). Unlike static methods, dynGENIE3 explicitly models temporal dependencies, making it well-suited for capturing dynamic regulatory interactions. dynGENIE3 is a machine learning-based method which employs ensemble decision trees as its core modelling technique. Decision trees are created to capture relationships between gene expression profiles, and an ensemble of these trees is used to enhance prediction accuracy. The algorithm selects a subset of relevant genes as candidate regulators for each target gene, where the functional relationships between target genes and their regulators are modelled with ordinary differential equation functions. The regulatory network is then built by associating each gene with a ranked list of potential regulators, representing the putative regulatory interactions between genes. The scores to candidate regulators are assigned based on their importance in predicting the expression of the target gene and indicating the strength of regulatory relationships.

The method takes as input (1) the TF activities calculated as above at each time point, (2) the indicated *time points* in which TF activity is measured, and (3) the *expression decay rates*, calculated directly from the input data. In our case, the dataset consists of the most differentially activated TFs in autologous culture and monocultures, for which we calculate TF-specific decay rates and use the median of the decay rates to run the algorithm for GRN inference. To generate the final dynGENIE3 network, we apply the method for 𝑛 = 10 times, and the intersection between the networks is considered for the subsequent analysis. In addition, we perform sensitivity and robustness analysis of the method’s parameters by exploring the tree-based method (Random Forest or Extra-Trees) and `ntrees`. The results show a high robustness of the method, quantified by taking the overlap between the inferred GRNs generated from each parameter choice.

#### TF differential activity correlation network

The differential TF activity is calculated between each successive time interval using msVIPER, producing a differential activity score that is set to 0 for all 𝑝*_val_* > 0. 05. For autologous and monoculture conditions separately, we consider a dataframe with values of differential activity in successive time intervals for all TFs that were selected previously as variables across time in the specific condition (see transcription factor activity estimation). We then calculate Pearson’s correlation of these differential activity values for all possible pairs of TFs and consider |0.5| as the weight threshold for establishing edges connecting the TFs.

Since this method did not provide a direction for the interaction, we consider the overlap of this TF differential activity correlation network with the dynGENIE3 inferred network to produce the final TF network shown in Figure 4 (a). Lastly, the module detection is performed using the ClusterViz Cytoscape plug-in (Wang et al., 2015) (FAG-EC algorithm, size threshold = 5) (Figure S.8 (b), Figure S.10). We identify 11 modules on the TF-TF network, on which we perform Over-Representation Analysis of their TFs and their target genes (Figure S.11).

### Data and code availability

The pre-processed data, the R scripts used for data pre-processing and analysis, gene regulatory network inference, as well as the session details (operating systems, RStudio version and libraries) and code to reproduce the figures, are available at https://github.com/VeraPancaldiLab/CLL_GRN_paper.

### Funding

This work was funded by the Agence Nationale de la Recherche (ANR-23-CE45-0014), granted to V.P. H.C. was funded by a CIFRE fellowship (No. 231967A10) established between V.P., A.Z., and H.C., in collaboration with Inserm Transfert, CNRS, Université Toulouse III – Paul Sabatier, the CHU de Toulouse, and EVOTEC (France) SAS. This study has been partially supported through the grant EUR CARe N°ANR-18-EURE-0003 in the framework of the Programme des Investissements d’Avenir, granted to M.H. The funders had no role in study design, data collection and analysis, decision to publish, or preparation of the manuscript.

## Acknowledgement.

The authors thank the members of the ANR RD2Bol consortium (Dr. Laurence Calzone, Dr. Loïc Paulevé and Victoria Bruning) for their valuable insights and feedback.

## Author contributions

Conceptualisation: M.M, H.C, V.P; Sample collection: L.Y; Experimental design and data collection: J.B, V.P.; Computational workflow M.M, H.C, V.P, A.Z; Data exploration: M.M, H.C, M.H, F.R, V.P.; Project supervision: M.M, V.P, A.Z; Interpretation of results: M.M, M.D, M.P, V.P, A.Z; Writing original draft: M.M, H.C, V.P., J.B; Review and editing: all authors.

## Supplementary Material

### Deconvolution analysis

To computationally calculate the cell percentages in bulk RNA-seq data of the autologous culture and disentangle cell–type–specific contributions within autologous and monoculture cultures, we perform deconvolution analysis using single-cell-derived signatures to estimate specific cell-type proportions from bulk RNAseq data as well as to infer the cell-type-specific expression profiles. Cell type signatures were generated with CIBERSORTx (CBSX) (Newman et al., 2015) and DWLS (Tsoucas et al., 2019) from scRNA-seq data obtained from 1 CLL patient PBMC, at three conditions: untreated, treated and at relapse (Cadot et al., 2020). Pseudobulking of the single-cell data was performed using row sums of the counts as the aggregation method, followed by TPM normalisation. From this, cell type deconvolution was applied using the CBSX and DWLS methods using the previously created signatures. In parallel, BayesPrism (Chu et al., 2022) method is also used and applied directly to the single-cell data, as it does not rely on precomputed static signatures. Method-signature performance is assessed using the ground truth proportions extracted from the single-cell data as reference and comparing them with the proportions estimated from the deconvolution using the mean squared error. Among the tested approaches, CBSX and DWLS-based signatures using the CBSX algorithm show the strongest performance (Figure S.1). Based on these results, cell deconvolution of bulk data from autologous and monoculture experiments is carried out using CBSX with the custom signatures. Results produced are coherent and biologically consistent with the FACS measurements. All the necessary files and R scripts to reproduce the results can be found in <u>Supplementary file</u>.

**Figure S.1:**
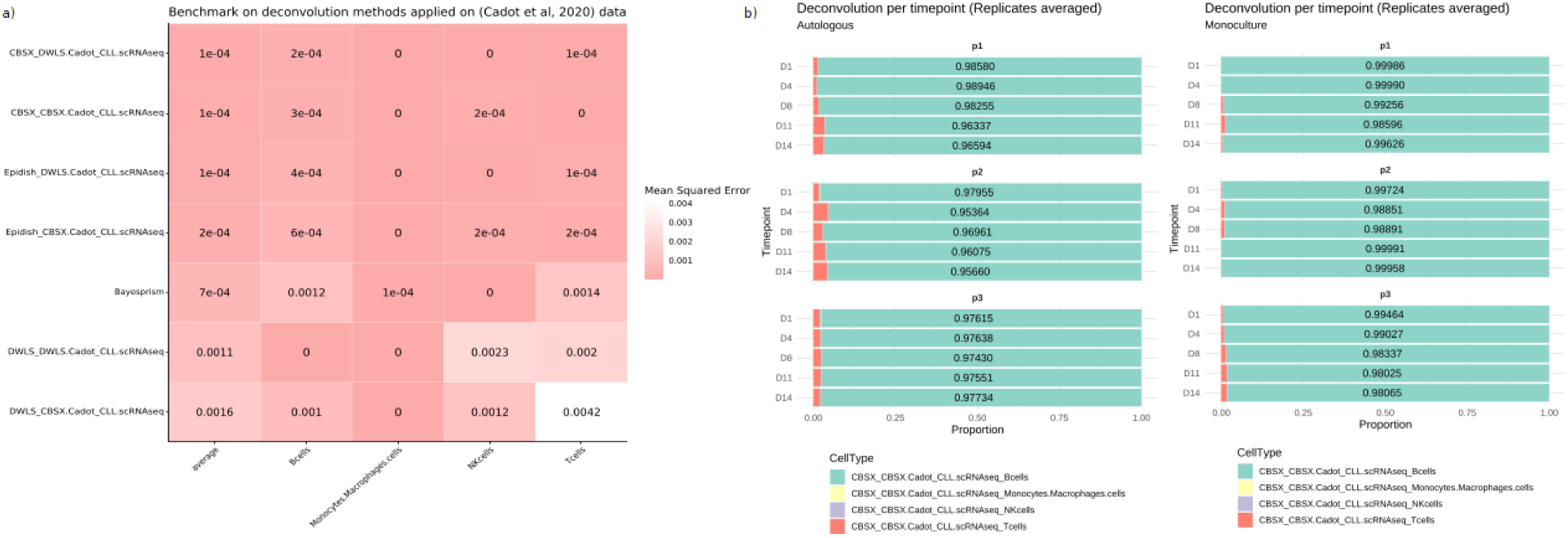
Deconvolution performed on autologous and monoculture, using single-cell RNAseq-derived signatures. (a) Heatmap with performance of signatures, (b) Cell percentages in autologous culture and monoculture, using the custom signatures with CBSX.

**Figure S.2:**
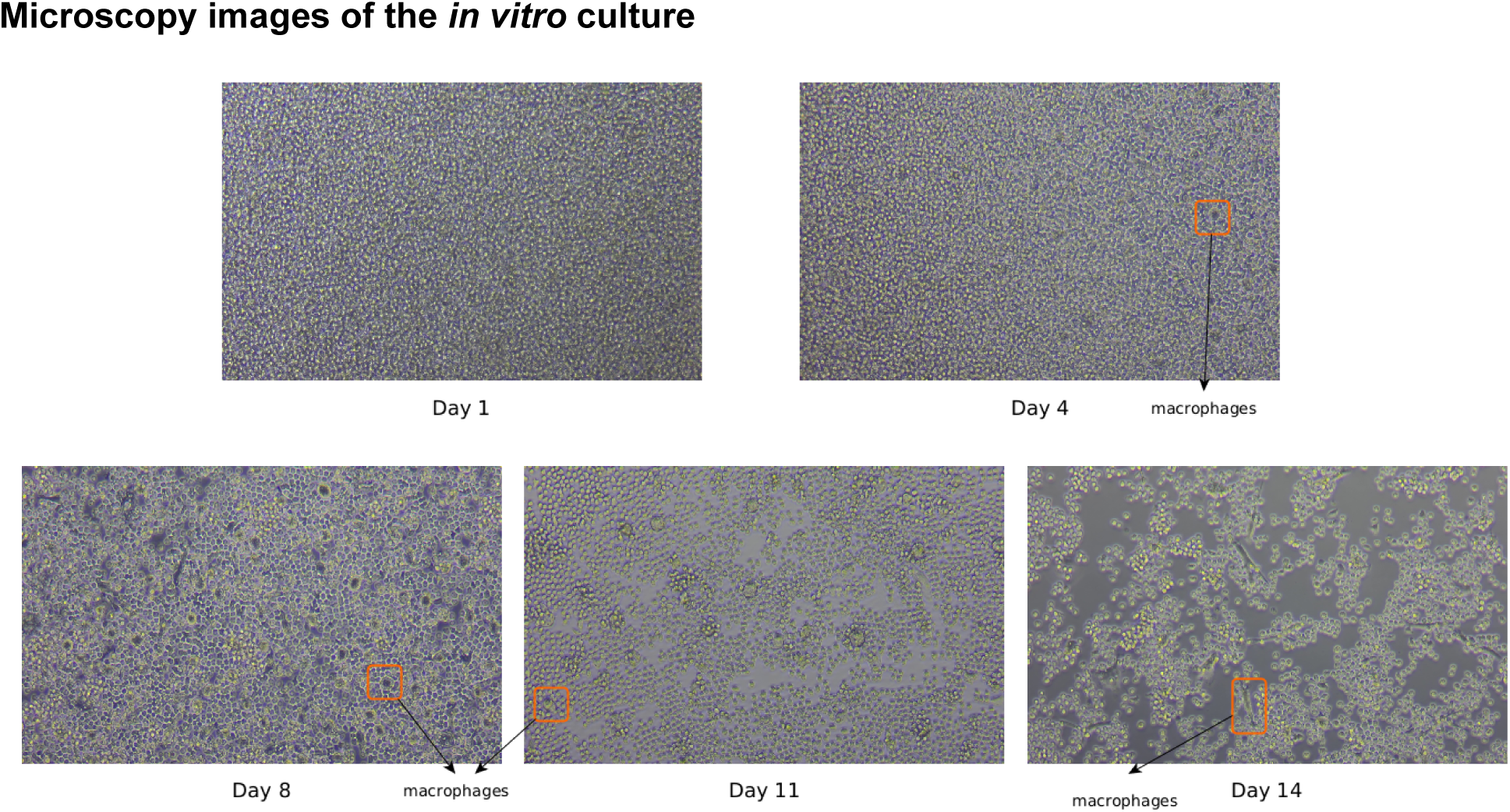
Microscopy images of the in vitro culture. After D8, the formation of macrophages is complete.

**Table S.1:**
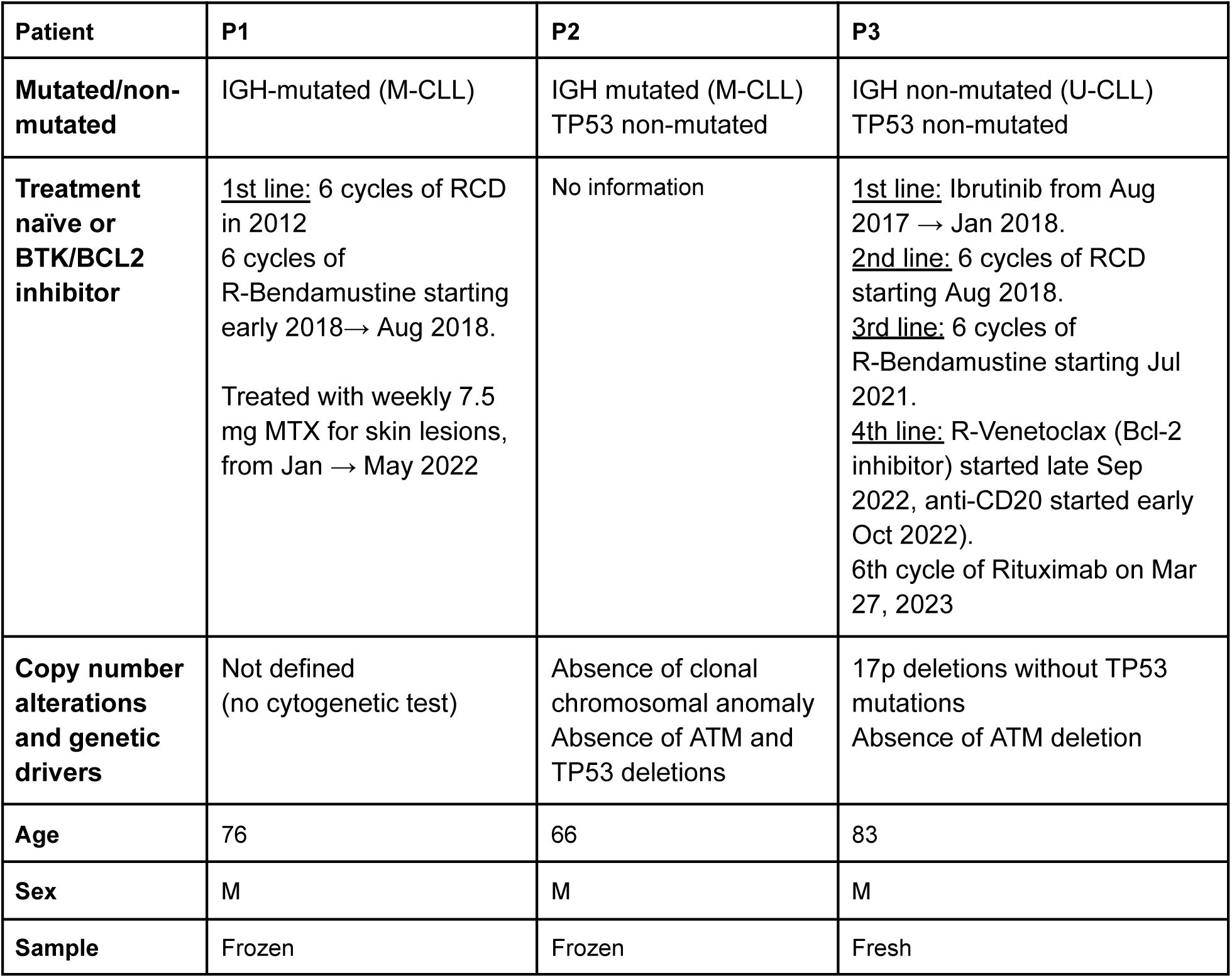
Clinical data of three CLL patients.

| Patient | P1 | P2 | P3 |
| --- | --- | --- | --- |
| **Mutated/non-mutated** | IGH-mutated (M-CLL) | IGH mutated (M-CLL)<br>TP53 non-mutated | IGH non-mutated (U-CLL)<br>TP53 non-mutated |
| **Treatment naïve or BTK/BCL2 inhibitor** | <u>1st line:</u> 6 cycles of RCD in 2012<br>6 cycles of R-Bendamustine starting early 2018→ Aug 2018.<br>Treated with weekly 7.5 mg MTX for skin lesions, from Jan → May 2022 | No information | <u>1st line:</u> Ibrutinib from Aug 2017 → Jan 2018.<br><u>2nd line:</u> 6 cycles of RCD starting Aug 2018.<br><u>3rd line:</u> 6 cycles of R-Bendamustine starting Jul 2021.<br><u>4th line:</u> R-Venetoclax (Bcl-2 inhibitor) started late Sep 2022, anti-CD20 started early Oct 2022).<br>6th cycle of Rituximab on Mar 27, 2023 |
| **Copy number alterations and genetic drivers** | Not defined (no cytogenetic test) | Absence of clonal chromosomal anomaly<br>Absence of ATM and TP53 deletions | 17p deletions without TP53 mutations<br>Absence of ATM deletion |
| **Age** | 76 | 66 | 83 |
| **Sex** | M | M | M |
| **Sample** | Frozen | Frozen | Fresh |

**Figure S.3:**
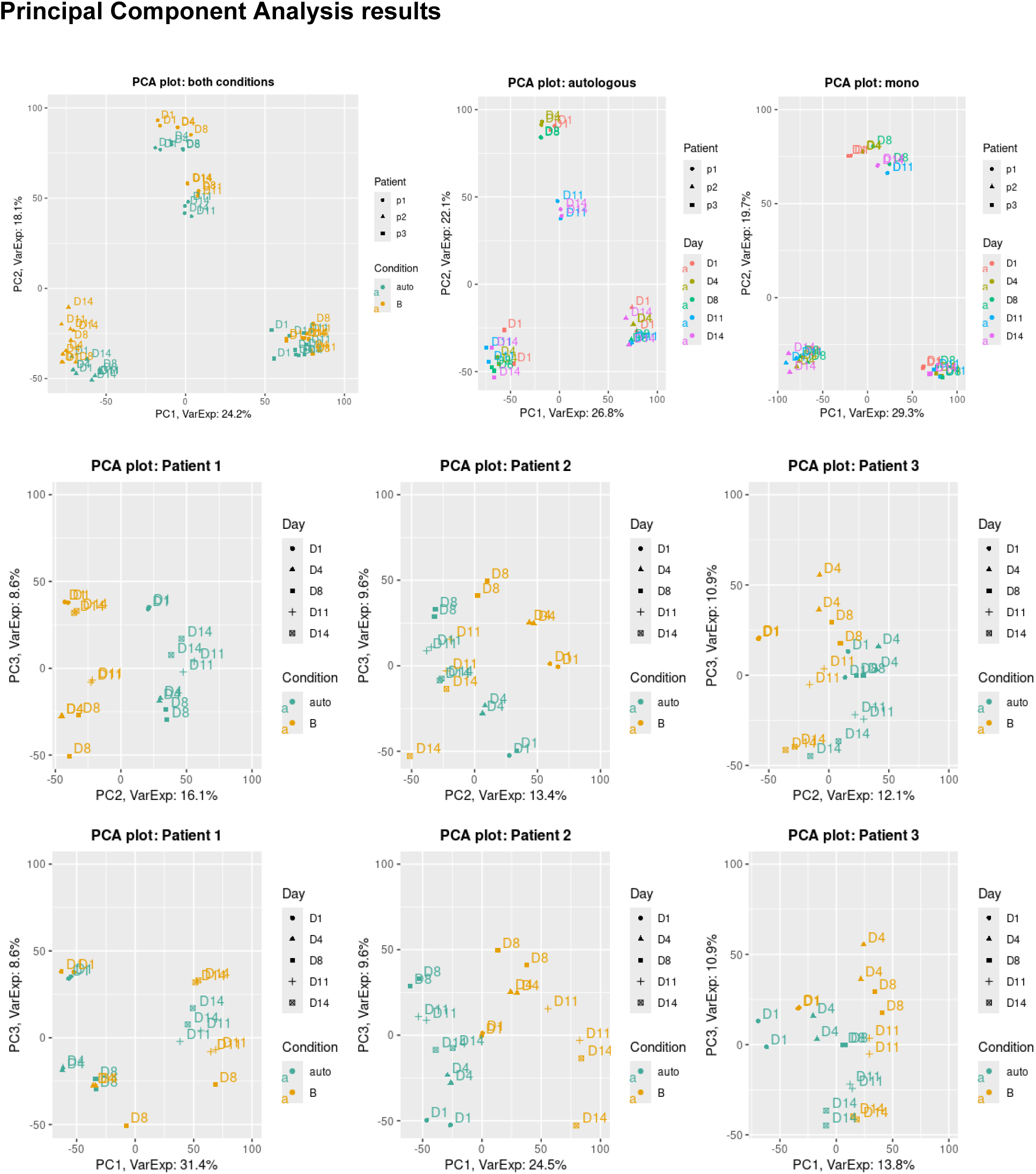
Principal Component Analysis of autologous and monoculture, showing the high heterogeneity between patients.

**Figure S.4:**
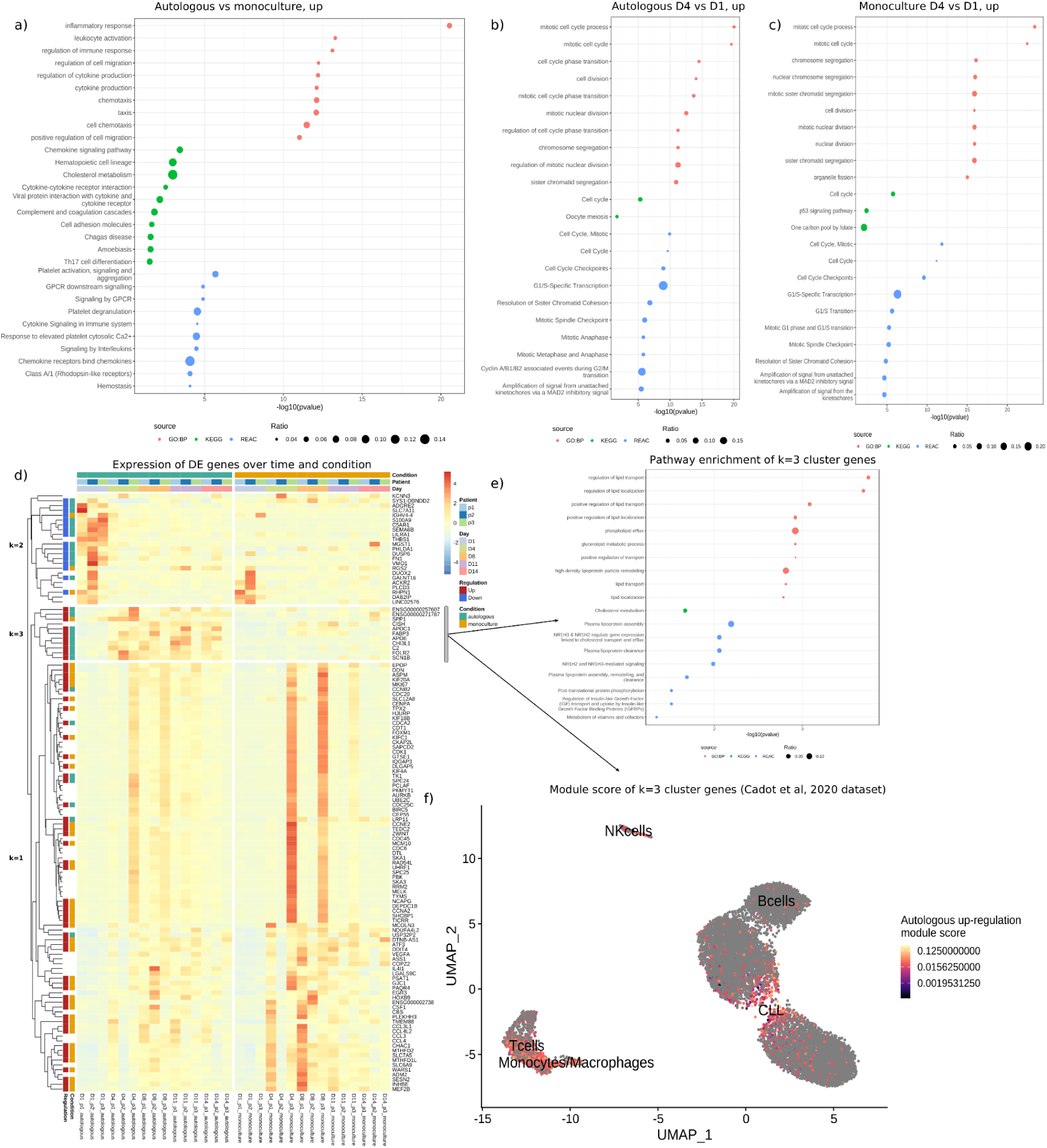
(a) Pathway enrichment analysis of the up-regulated genes in the autologous vs monoculture. (b), (c) Pathway enrichment analysis of the up-regulated genes in D4 vs D1 comparison in the autologous and monoculture, respectively. (d) Expression of the differentially expressed genes in autologous and monoculture in 𝑡 𝑣𝑠 𝑡 − 1 comparison. (e) Pathway enrichment analysis of the 𝑘 = 3 genes. (f) UMAP of CITE-seq dataset of CLL PBMC from (Cadot et al, 2020) and score of the 𝑘 = 3 gene module.

**Figure S.5:**
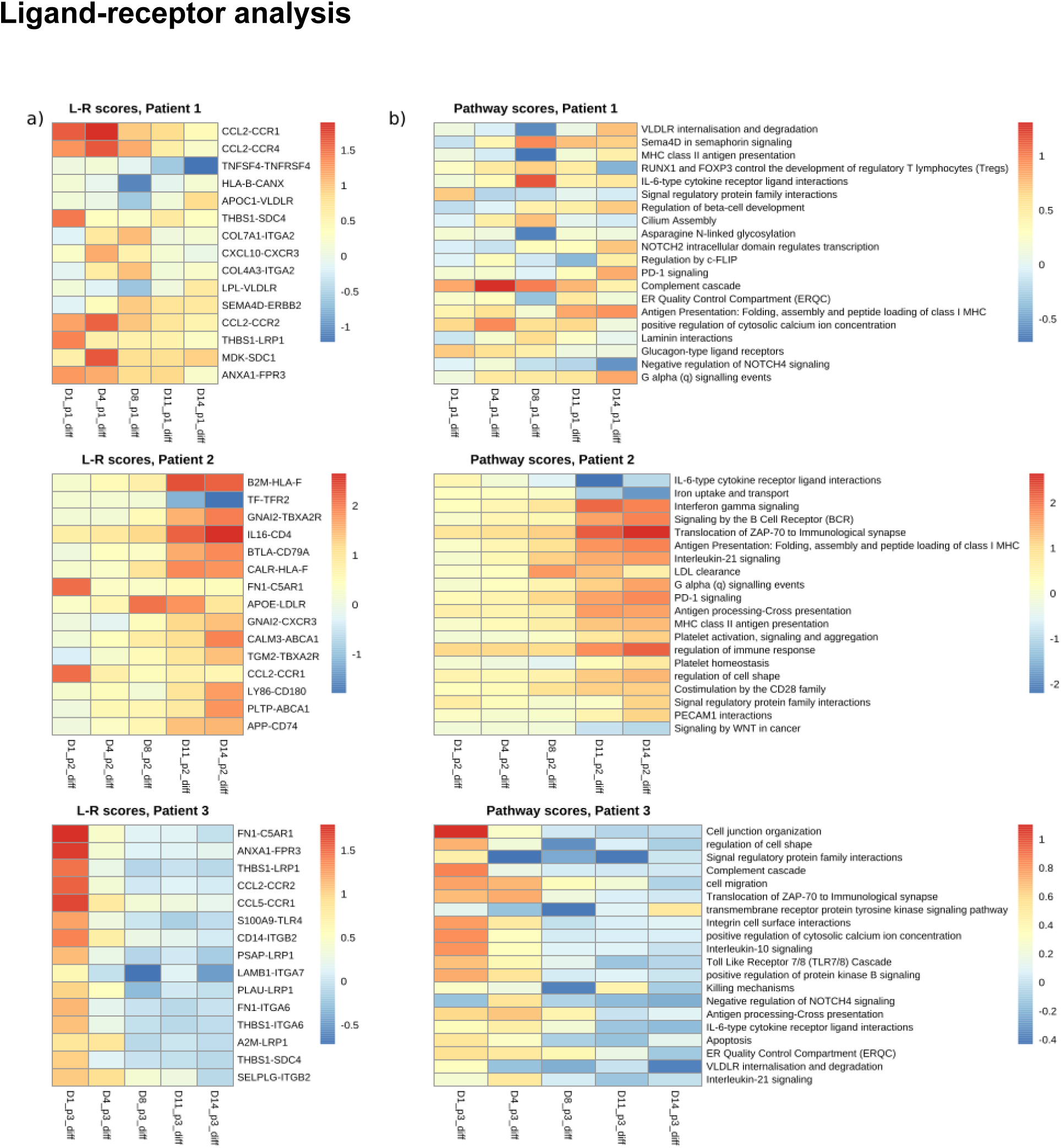
(a) Ligand-receptor pair activation. (b) Pathway scores for the expressed L-R pairs.

**Figure S.6:**
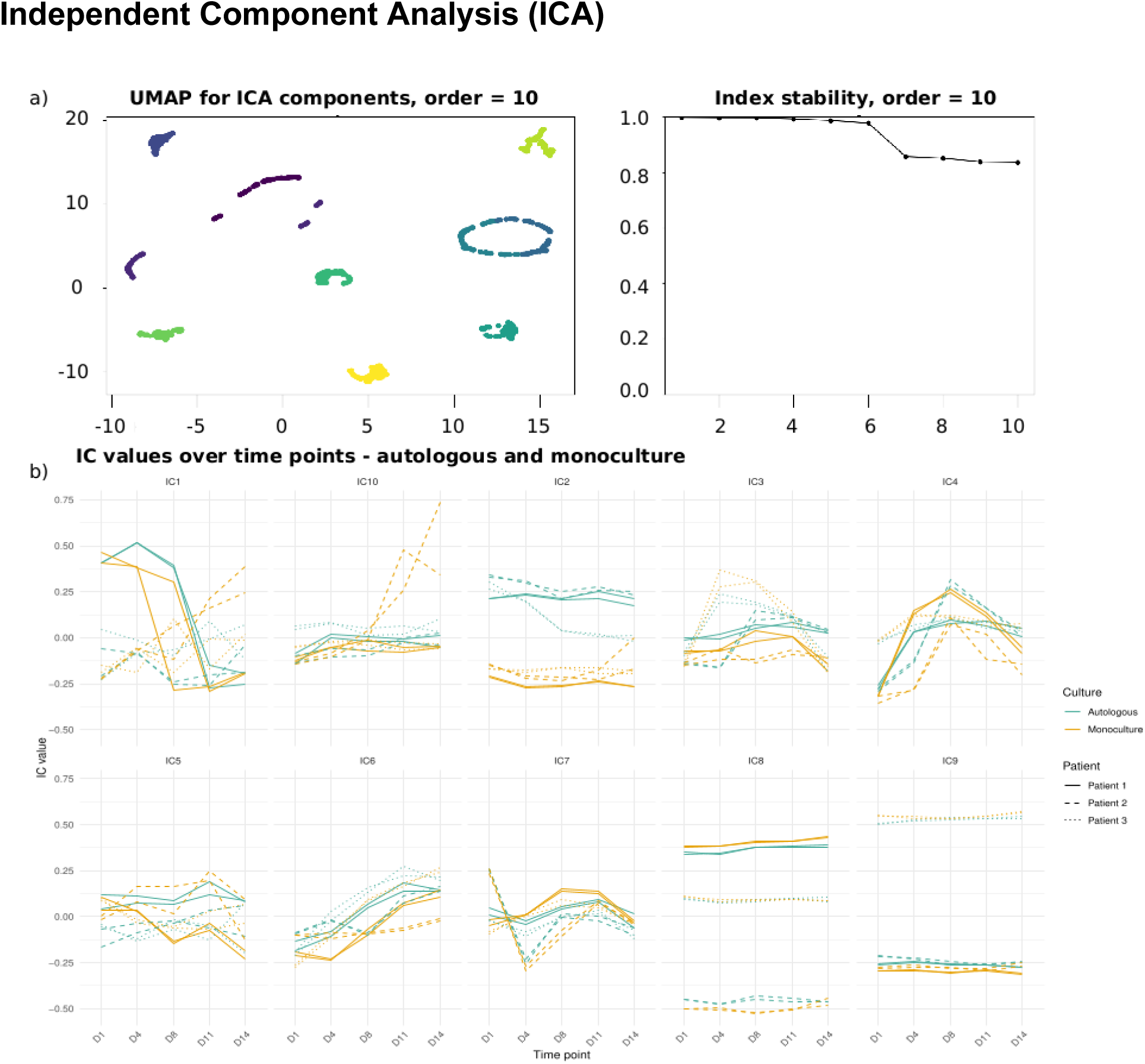
**(a)** Index stability - autologous and monoculture. **(b)** Heatmap of the metasample matrix from ICA. **(c)** ICA - TF activity score.

**Figure S.7:**
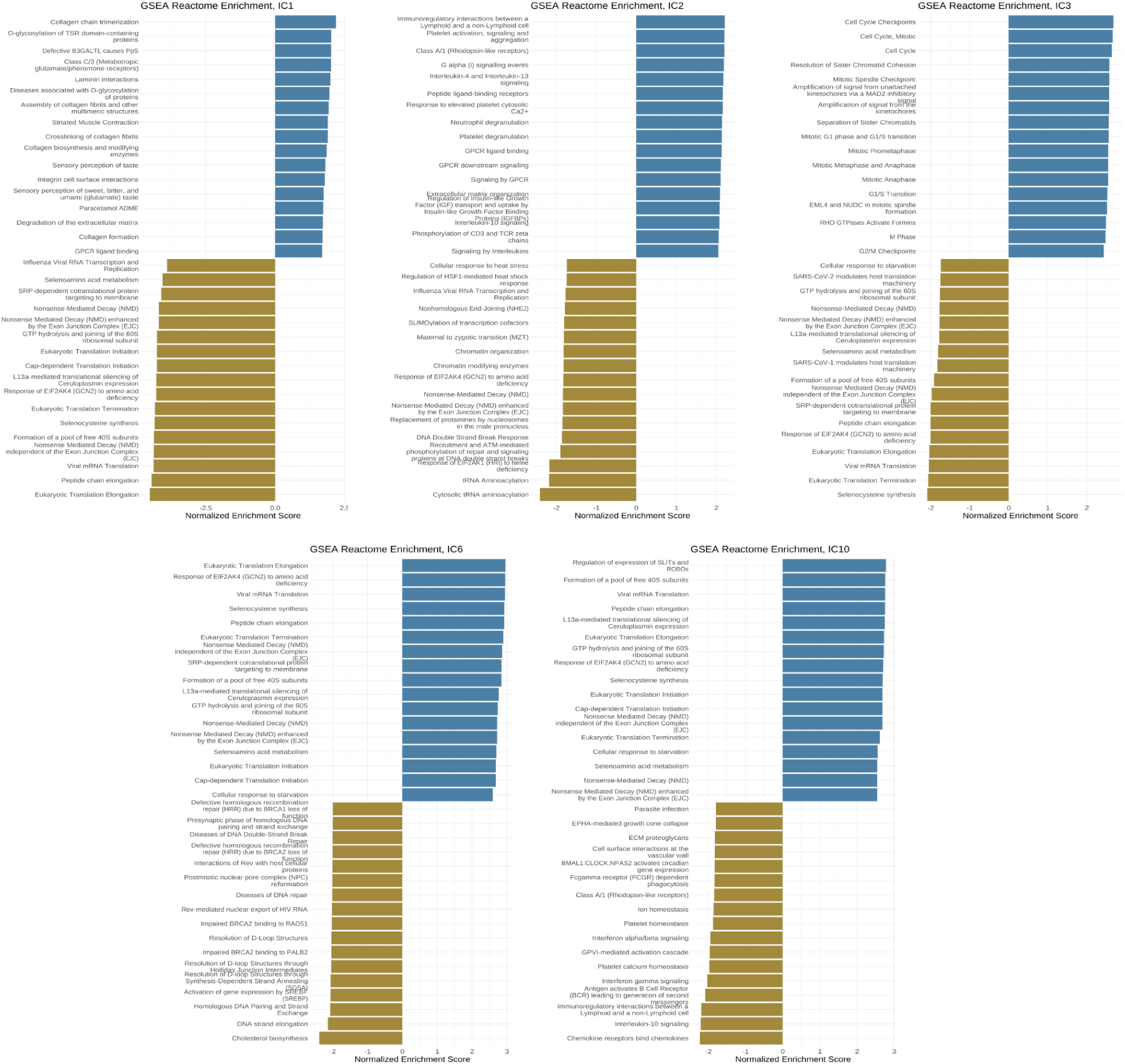
Gene set enrichment analysis on the independent components.

**Figure S.8:**
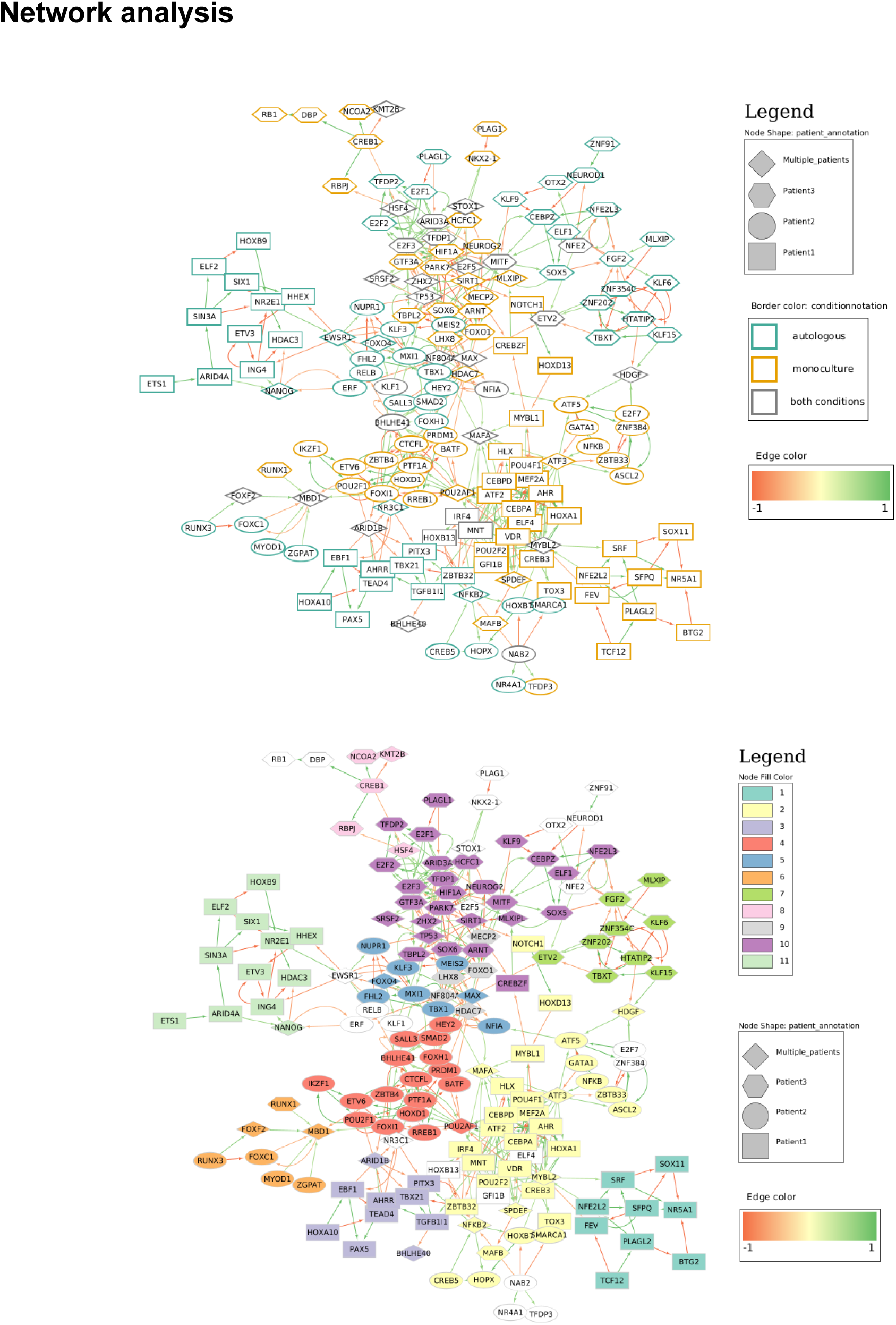
(a) TF-TF network with nodes coloured by condition. (b) TF-TF network with nodes coloured by module.

### Validation of TF selection through external dataset analysis

To further evaluate the robustness of our transcription factor (TF) selection strategy, we perform an external validation using single-cell RNA-seq data from CLL samples published by (Cadot et al., 2020). For this analysis, we employ the pySCENIC framework (Aibar et al., 2017; Kumar et al., 2021), which integrates regulatory network inference with cis-regulatory motif enrichment to prune false-positive edges and subsequently identifies master regulators capable of distinguishing transcriptional sub-populations.

Applied to the CLL population in the Cadot et al. dataset, pySCENIC identifies 284 master TFs that discriminate between pre-treatment and post-treatment states. We then compare these regulators with the set of TFs retained in our inferred GRN (167 TFs in total), obtaining an overlap of 59 TFs. Considering the differences in patient cohorts, timing of sequencing, and experimental protocols, we regard this degree of concordance as significant, supporting the notion that our TF selection approach successfully prioritises regulators central to the dynamic behaviour of CLL cells.

In addition, mapping the overlapping TFs onto our inferred GRN reveal that they predominantly occupy central positions in the network and are particularly enriched at the earliest time points of the culture (Figure S.11). These observations suggest that the common TFs capture core biological processes of CLL cells, which appear conserved across both the *in vitro* autologous culture and monoculture cultures and the independent CLL patient cohort profiled by Cadot et al.

All the necessary scripts to reproduce the results can be found in <u>Supplementary file</u>.

**Figure S.9:**
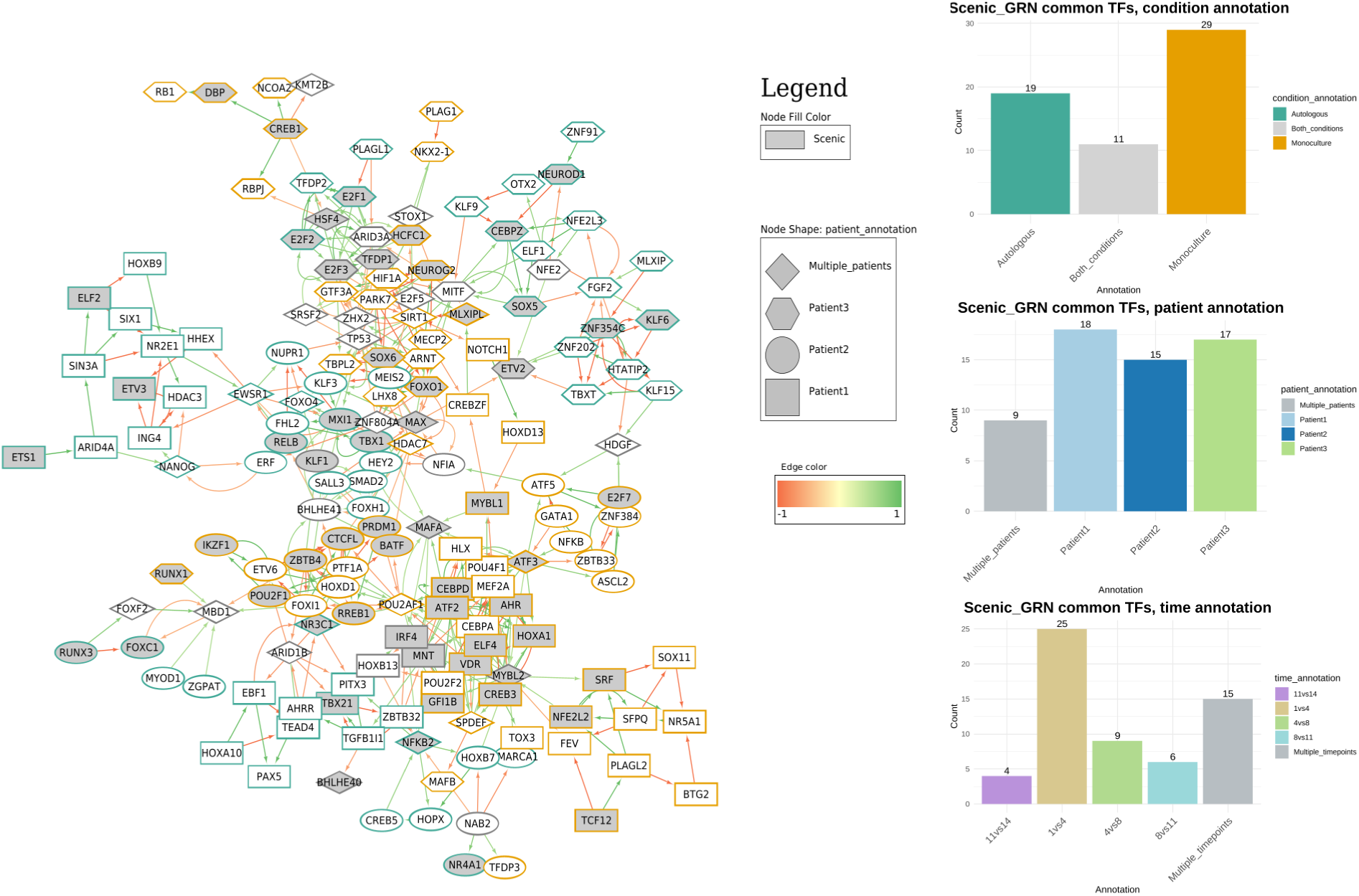
**(a)** Mapping of the overlapping TFs onto our inferred GRN. **(b)** Partition of the overlapping TFs according to condition, patient and time-point.

**Figure S.10:**
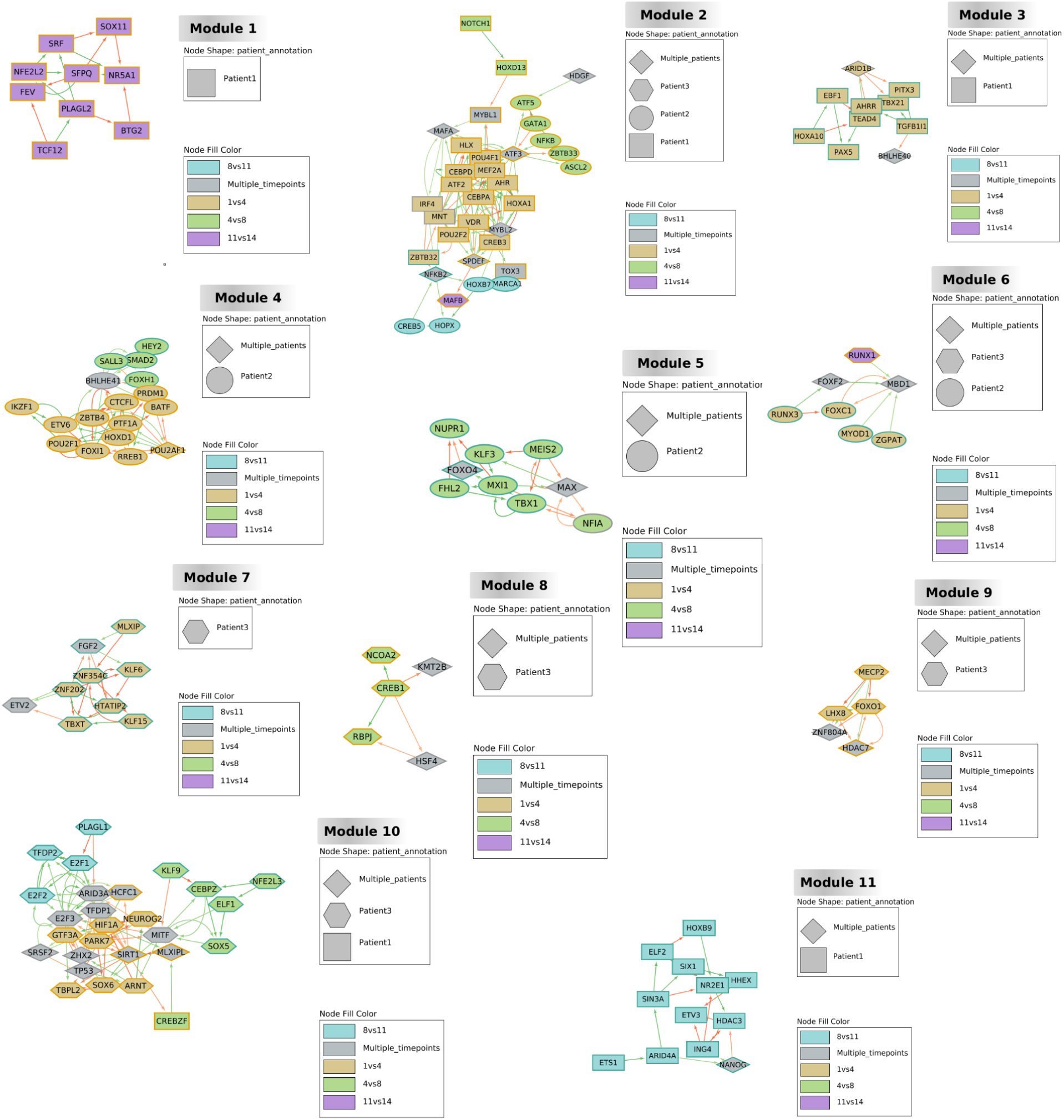
Annotation of modules with nodes’ features.

**Figure S.11:**
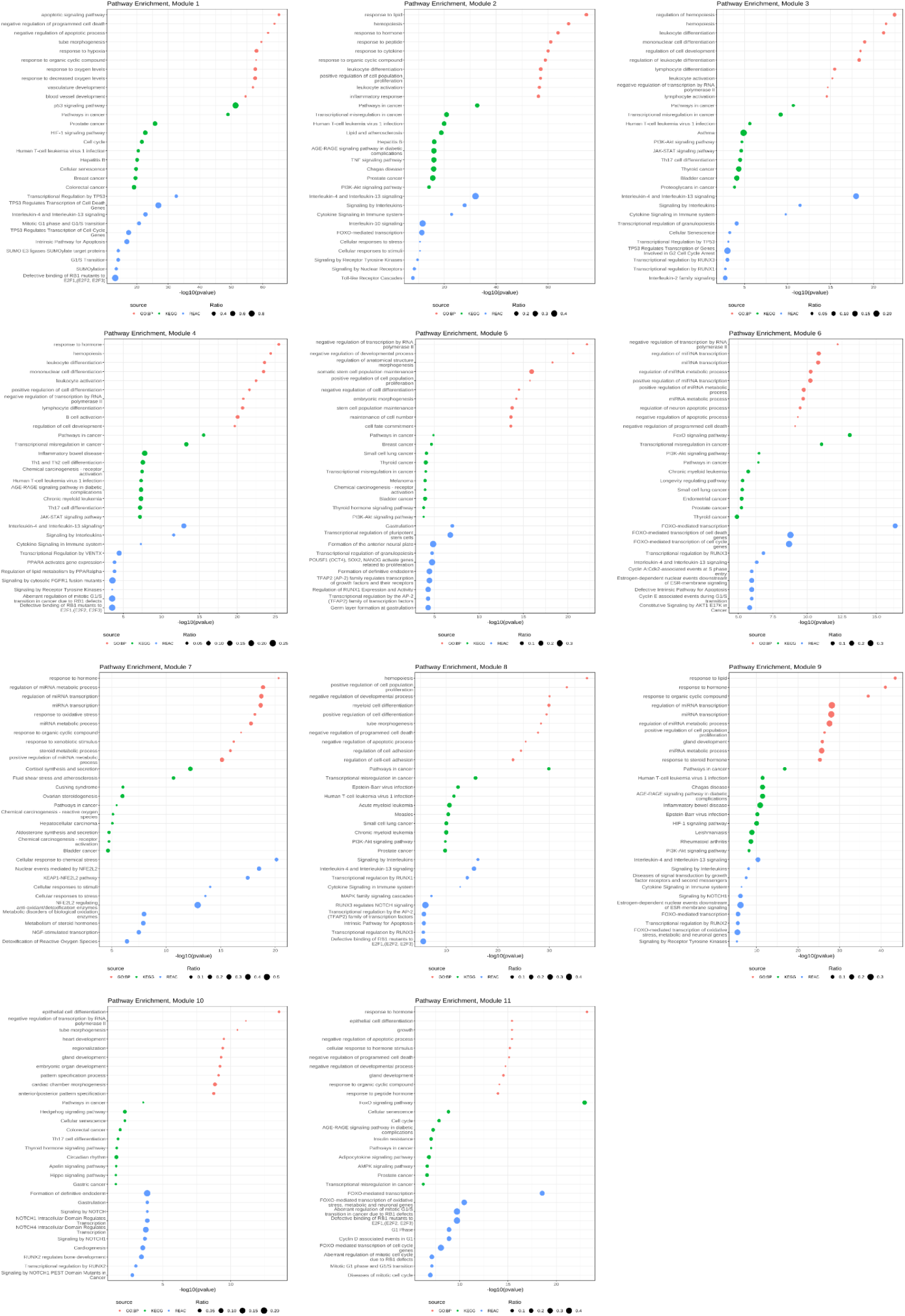
Pathway enrichment analysis of each cluster of TFs (including their targets).

## Notes

https://github.com/VeraPancaldiLab/CLL_GRN_paper

### Competing Interest Statement

The authors have declared no competing interest.

### Summary of Updates

This version of the manuscript has been revised to update the following: (1) deconvolution analysis, (2) network inference and validation with external datasets.

